# LARP7 is a BRCA1 ubiquitinase substrate and regulates genome stability and tumorigenesis

**DOI:** 10.1101/808535

**Authors:** Fang Zhang, Pengyi Yan, Huijing Yu, Huangying Le, Zixuan li, Jiahuan Chen, Xiaodong Liang, Shiyan Wang, Weiting Wei, Li Liu, Yan Zhang, Xing Ji, Anyong Xie, Wantao Chen, Zeguang Han, William T. Pu, Sun Chen, Yingwei Chen, Kun Sun, Baoxie Ge, Bing Zhang

## Abstract

Attenuated DNA repair leads to genomic instability and tumorigenesis. BRCA1/BARD1 are the best known tumor suppressors that promote homology recombination (HR) and arrest cell cycle at G2/M checkpoint. As E3 ubiquitin ligases, their ubiquitinase activity has been known to involve in the HR and tumor suppression, but the mechanism remains ambiguous. Here, we demonstrated upon genotoxic stress, BRCA1 together with BARD1 catalyzed the K48 ployubiquitination on LARP7, a 7SK RNA binding protein known to control RNAPII pausing, and thereby degraded it through 26S ubiquitin-proteasome pathway. Depleting LARP7 suppressed the expression of CDK1 complex, arrested cell at G2/M DNA damage checkpoint and reduced BRCA2 phosphorylation which thereby facilitated RAD51 recruitment to damaged DNA to enhance HR. Importantly, LARP7 depletion observed in breast patients lead to the chemoradiotherapy resistance both *in vitro and in vivo*. Together, this study unveils a mechanism by which BRCA1/BARD1 utilizes their E3 ligase activity to control HR and cell cycle, and highlights LARP7 as a potential target for cancer prevention and therapy.

**Highlights:**

1. DNA damage response downregulates LARP7 through BRCA1/BARD1
2. BRCA1/BARD1 catalyzes the K48 polyubiquitination on LARP7
3. LARP7 promotes G2/M cell cycle transition and tumorigenesis via CDK1 complex
4. LARP7 disputes homology-directed repair that leads to tumor therapy resistance

## Introduction

A double-strand break (DSB) is the most deleterious type of DNA damage. The aberrant repair of DSB can cause genome instability, cell death, or uncontrolled cell growth (Ciccia and Elledge, 2010). There are two main mechanisms to repair DSBs: the error-free repair pathway of homology-directed recombination (HDR) and the error-prone repair pathway of non-homologous end joining (NHEJ).

Breast cancer type I susceptibility protein (BRCA1) is a tumor suppressor that is mutated in 3-18% of all breast and ovarian cancer and over 45% of hereditary breast and ovarian cancer (Miki et al., 1994) (Huen et al., 2010). BRCA1 mutation has emerged as a genetic biomarker for the clinical diagnosis of breast and ovarian cancer(Miki et al., 1994) (Brody and Biesecker, 1998). BRCA1 is a ubiquitin E3 ligase that contains an N-terminal RING motif and two BRCA1 C terminus (BRCT) motifs. BRCA1 suppressed tumor progression primarily by controlling the G2/M checkpoint and enhancing HDR(Huen et al., 2010; Moynahan et al., 1999). Sites of DNA damage are marked by ubiquitinated histones, which bind the ubiquitin-interacting motif (UIM) of RAP80, which in turn recruits BRCA1 (Kim et al., 2007; Wang et al., 2007). Chromatin-bound BRCA1 further recruits the CTBP-interacting protein (CtIP) and meitotic recombination 11 (MRE11)-RAD50-NBS1Complex (MRN complex) and resected the damaged DNA ends(Chen et al., 2008; Yun and Hiom, 2009). Moreover, BRCA1 enhanced homologous recombination by increasing PALB2-BRCA2-RAD51 recruitment to DNA damage foci and by activating RAD51-mediated strand invasion and D-loop formation(Sy et al., 2009; Zhao et al., 2017).

Beyond facilitating the formation of complexes involved in HDR, BRCA1 ubiquitin E3 ligase activity has been long suspected to have an irreplaceable role in DNA repair and tumor suppression (Morris and Solomon, 2004). Numerous disease-causing mutations in RING domain of BRCA1 and its major partner BRCA1-associated RING domain protein 1 (BARD1, also a RING domain E3 ligase) were discovered in hereditary or early-onset breast and ovarian cancer (Thai et al., 1998). BRCA1 dimerizes with BARD1 via their RING domains and ligates the non-canonical ubiquitin linkage (K6 and K63) to H2A, H2A.X and CtIP (Yu et al., 2006) (Chen et al., 2002). Although essential for DNA damage repair, the mechanism by which the BRCA1/BARD1 ubiquitin ligase promotes DNA damage repair remains unclearly defined.

La-related protein7 (LARP7) is a La family RNA-binding protein containing two RNA binding domains: RNA recognition motif (RRM) and HTH La-type RNA-binding domains (Krueger et al., 2008) (Markert et al., 2008). LARP7 binds to and stabilizes the 3’ hairpin of 7SK RNA, the most abundant noncoding RNA in mammalian cells, to form the core of the 7SK snRNP (7SK small nuclear ribonucleoprotein). 7SK snRNP thus far has been identified to negatively regulate RNAPII pausing release, a critic RNA transcription checkpoint widespread in many organisms, by sequestering the positive transcription elongation factors pTEFb complex in the nucleoplasm (Yang et al., 2001) (Nguyen et al., 2001). The attenuation of LARP7 disrupts 7SK snRNP, relocates pTEFb to chromatin and therefore releases RNAPII pausing. Recessive *LARP7* loss-of-function mutations were discovered to cause primordial dwarfism in humans and mice, possibly by suppressing cell growth(Alazami et al., 2012). Moreover, LARP7 has been reported to suppress the tumorigenesis of gastric cancer and cancer progression and metastasis in breast cancer as a tumor suppressor(Cheng et al., 2012; Ji et al., 2014). However, the functions of LARP7 and 7SK snRNP beyond the RNAPII pausing such as in the DDR and DNA damage repair remain unknown.

In this study, we identified LARP7 as a novel K48 ubiquitination substrate of BRCA1/BARD1. BRCA1/BARD1 ubiquitination of LARP7 was required for BRCA1 control of the G2/M checkpoint and HDR, and for its activity as a tumor suppressor. This discovery provides answers for the long standing question of whether and how BRCA1 uses it’s E3 ligase activity to maintain the genome stability and suppress tumorigenesis, and unveils a new DDR pathway centered on 7SK snRNP.

## Results

### LARP7 deficiency attenuated the DNA damage response

Ionizing radiation (IR) induces a profound DNA damage response (DDR) mediated by ATM, a kinase activated by self-phosphorylation (Ciccia and Elledge, 2010). A previous proteomics study revealed that LARP7 is a potent kinase substrate of ATM and indicated that it might be involved in the DDR (Matsuoka et al., 2007). To test this possibility, we examined LARP7 expression in IR-treated mouse embryonic fibroblasts (MEFs) and found that IR caused a gradual reduction in LARP7 along with DDR activation, as indicated by ATM phosphorylation and BRCA1 upregulation (Fig. 1a). LARP7 is a 7SK RNA binding protein to maintain the 7SK snRNP stability (Muniz et al., 2013). Lost of LARP7 in IR-treated MEF disrupted the 7SK RNP as illustrated by a decrease of 7SK RNA (sFig. 1a). Degradation of LARP7 were also identified through immunofluorescence staining with a LARP7-specific antibody (Fig. 1b). Intriguingly, an extranuclear shuttling of LARP7, after IR, was also observed, which further affirmed a collapse of 7SK snRNP upon IR because the intact 7SK snRNP prominently lodges within the nucleus (Fig. 1b). Cisplatin (CDDP), a widely used first-line chemotherapy drug for cancer, activates the DDR and causes DNA damage mainly by generating DNA crosslinks. Consistent with the results of IR, CDDP upregulated BRCA1, phosphorylated ATR and ATM (DNA damage sensors), and induced significant LARP7 and 7SK RNA downregulation and extranuclear shuttling in MEFs (sFig. 1b-d). The downregulation of LARP7 was also observed in CDDP-treated HeLa and further confirmed with a second, independent LARP7 antibody (sFig. 1e,f).

**Figure 1.**
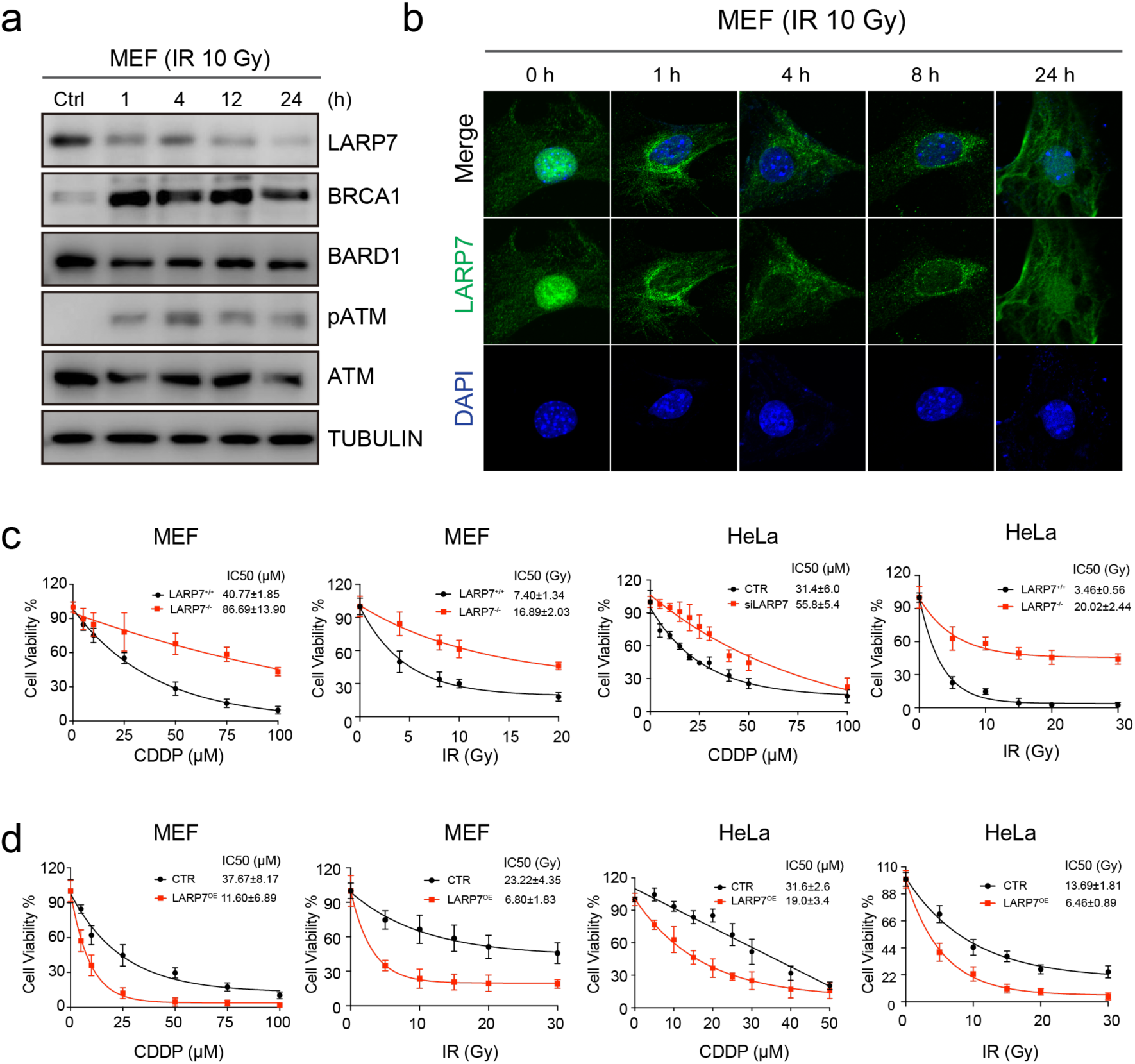
LARP7 dampened the DDR. a, WB showing LARP7 degradation during the IR-induced DDR. b, LARP7 immunofluorescence staining showing that LARP7 was shuttled out of nuclei and was diminished after IR treatment. c-d, LARP7 deletion led to IR and CDDP resistance in MEFs and HeLa cells (c); conversely, forced LARP7 upregulation increased cell sensitivity. (d). Cell viability was assessed with a CCK8 assay. Data: mean±SD, regression: one-phase decay model. The IC50 was determined with a one-phase decay model.

Excessive DDR and DNA damage can initiate cell death. To further implicate the involvement of LARP7 in the DDR, we compared the survival of LARP7-knockout or LARP7-knockdown MEFs and HeLa cells to that of control cells after DDR activation. Suppression of LARP7 in MEFs and HeLa cells prevented cell death induced by both IR and CDDP, as assessed with a cell viability assay (Fig. 1c, sFig. 2a-d). In contrast, ectopic upregulation of LARP7 decreased the number of surviving cells (Fig. 1d). These results were further confirmed by clonogenic assays, which are not dependent on measurements of enzyme activity (sFig. 3a-c). An enhanced DNA repair increases cell survival by preventing lethal genomic instability. Thus, these results indicated that upon DDR activation, LARP7 degradation and resulted 7SK snRNP disassembly might prevent genotoxic stress induced genome instability via an enhanced DNA repair.

### The BRCA1/BARD1 complex elicited the K48-linked polyubiquitination and degradation of LARP7

We first set out to understand the mechanism underlying LARP7 downregulation in the DDR. We analyzed LARP7 transcription in MEFs and HeLa cells by RT-PCR and found no obvious transcript downregulation after IR (sFig. 4a-b), which indicated that a posttranscriptional mechanism regulates LARP7 degradation. We thus assessed LARP7 protein stability by measuring its half-life and observed faster LARP7 degradation rate in IR-treated HeLa cells over untreated cells (Fig. 2a). By ubiquitination assay, an increased polyubiquitination rather monoubiquitination on LARP7 was observed in IR-treated 293T cells (Fig. 2b).

**Figure 2.**
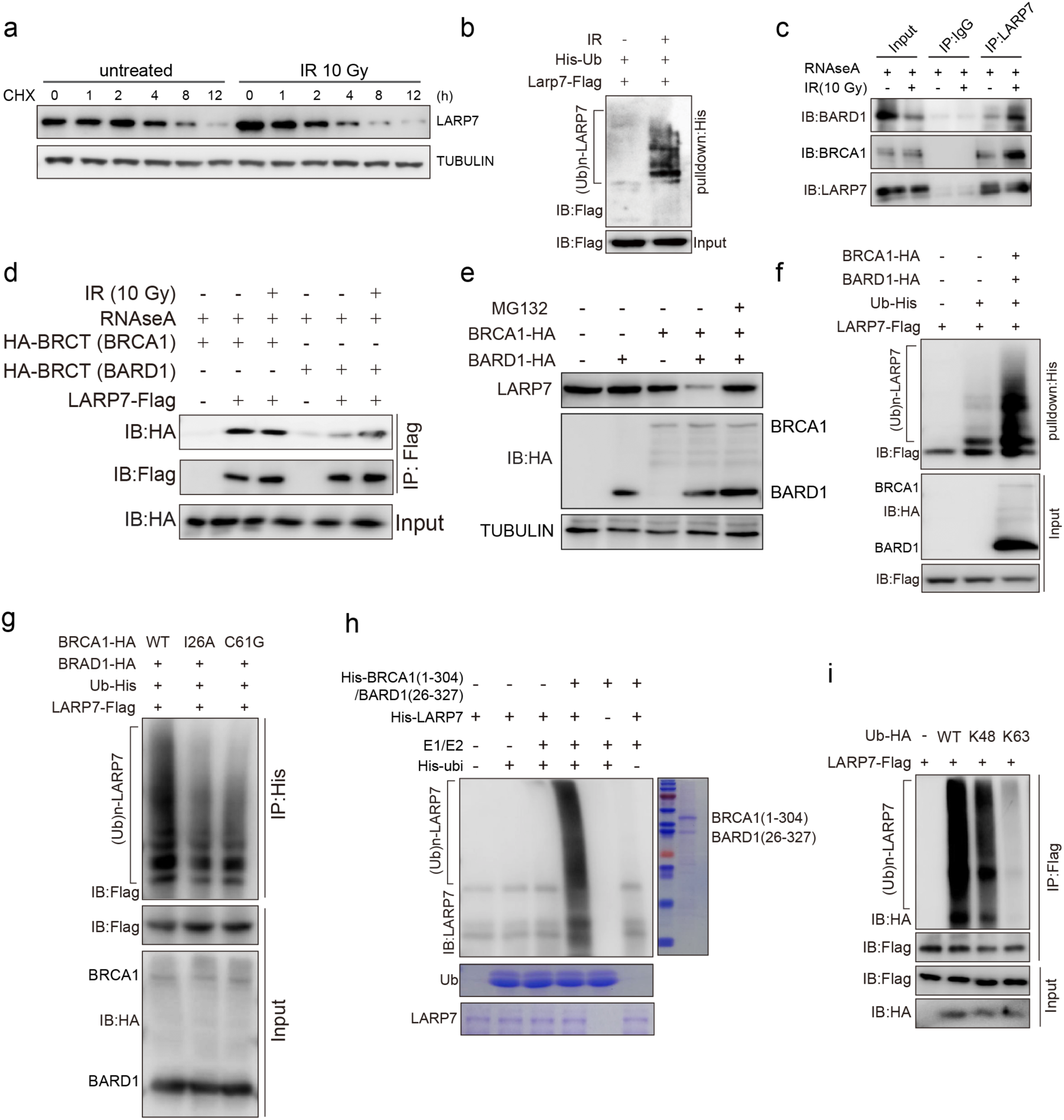
BRCA1/BARD1 catalyzed the K48 polyubiquitination on LARP7. a, IR reduced the half-life of LARP7. b, *In vivo* ubiquitination assay results showing that IR induced the ubiquitination of LARP7. c, LARP7 interacted with BRCA1 and BARD1, which was independent of 7SK RNA and was increased by IR. d, IR enhanced the interaction of LARP7 with the BARD1 BRCT domain but not the BRCA1 BRCT domain. e, Ectopic expression of BRCA1/BARD1 promoted LARP7 degradation, an effect that was attenuated by ubiquitination inhibition (MG132). f, BRCA1/BARD1 coexpression induced LARP7 ubiquitination in 293T cells. g. Loss-of-function mutations in the BRCA1 RING domain (I26A and C61G) abolished LARP7 ubiquitination. h. An *in vitro* ubiquitination assay illustrated that LARP7 ubiquitination was mediated by the RING domain complex of BRCA1 (1-304) and BARD1 (26-327). i. An *in vivo* ubiquitination assay demonstrated that K48-linked polyubiquitin was the predominant form of BRCA1-elicited LARP7 ubiquitination. K48: ectopically expressed ubiquitin with all lysine residues mutated to arginine except K48. K63: ubiquitin with all lysine residues mutated to arginine except K63. WT: wild-type ubiquitin with all lysine residues intact.

The specificity of protein ubiquitination is largely determined by E3 ligases. To determine the E3 ligase responsible for LARP7 ubiquitination, we mined the Biological General Repository for Interaction Datasets (BioGRID) protein interaction database and found that the E3 ligase BARD1 has been reported to interact with LARP7(Woods et al., 2012). BARD1 is a RING domain E3 ligase that confers ubiquitination after dimerizing with BRCA1, a RING domain E3 ligase as well (Xia et al., 2003) (Eakin et al., 2007). Multiple loss-of-function mutations within their RING domain identified in breast and ovarian cancers indicts the indispensable role of their E3 ligase activity in DNA repair and tumorigenesis. However, at present, no substrate directly regulating DNA repair has been identified (Elia and Elledge, 2012). To test if LARP7 could be the unidentified substrate for BRCA1/BARD1, we further validated this interaction in 293T cells by endogenous immunoprecipitation (Fig. 2c, sFig. 4c-d) and coimmunoprecipitation (Co-IP) (sFig. 4e). Both BARD1 and BRCA1 interacted with LARP7, and the interaction was augmented by IR and independent of 7SK RNA which was further confirmed by biomolecular fluorescence complementation (BiFC) assay in living cells (sFig. 4f-g).

The BRCT domain of BRCA1 and BARD1, the major region involved in interactions with other proteins, was sufficient for LARP7 interaction in the co-IP assay (Fig. 2d), consistent with the BioGRID data showing that the BRCT domain of BARD1 directly interacts with LARP7. IR increased LARP7’s interaction with both full-length BARD1 and BARD1 BRCT, but only increased the interaction with full-length BRCA1 not BRCA1 BRCT, which suggested that the enhanced interaction with full-length BRCA1 was passed from BARD1 that normally dimerizes with BRCA1 with their RING domains. Furthermore, these results implicated a posttranscriptional modification of LARP7 might modulate the interaction of LARP7 with BARD1 rather BRCA1 BRCT domain. This assumption was further supported by a GST pulldown assay showing that an unmodified LARP7 had a weak interaction with BARD1 BRCT comparing to BRCA1 BRCT(sFig. 4h).

To further interrogate if BRCA1/BARD1 is the ubiquitinase mediated LARP7 degradation, we ectopically expressed BRCA1/BARD1 in 293T cells and observed that BRCA1/BARD1 overexpression markedly decreased LARP7 expression; this effect was reversed by MG132, an inhibitor of the 26S proteasome (Fig. 2e). Conversely, knockdown of BRCA1 and BARD1 inhibited the LARP7 degradation induced by IR (sFig. 5a-b). In parallel, LARP7 expression in BRCA1 mutant cell lines of HCC1937 and MDA-MB-436 (frameshift mutations in the BRCT domain), was higher than that in BRCA1 wild-type cell lines (sFig. 5c). Furthermore, mutated BRCA1 significantly attenuated IR-induced LARP7 downregulation (sFig. 5d). Pulldown with either histidine (His)-tagged ubiquitin or with Flag-tagged LARP7 demonstrated ectopic expression of BRCA1 or both BRCA1/BARD1 increased LARP7 polyubiquitination in 293T cells (Fig. 2f, sFig. 5e). These lines of evidence together illustrated that BRCA1/BARD1 in fact ubiquitinated and degraded LARP7 during the DNA damage.

We further examine if the RING domain of BRCA1/BARD1 catalyze the LARP7 ubiquitination using two breast and ovarian cancer BRCA1 RING domain mutants, I26A and C61G, that abolish BRCA1 ubiquitination activity and are relevant to breast and ovarian cancer, and found that two mutations dramatically reduced LARP7 ubiquitination (Fig. 2g) (Hashizume et al., 2001) (Brzovic et al., 2003). An *in vitro* ubiquitination assay demonstrated that purified BRCA1 and BARD1 RING domain, constituted with E1 and E2 protein efficiently transferred the polyubiquitin chain to LARP7 (Fig. 2h), but either BRCA1 or BARD1 can’t complete this ubiquitination. By leveraging *in vivo* ubiquitination assay with transfection of K48- or K63-preserved ubiquitin, we further interrogated the polyubiquitination pattern of LARP7 upon IR induction. The results demonstrated K48-linked rather K63-linked polyubiquitin was introduced to LARP7 by IR, which further supported it is 26S proteasomal pathway that mediated LARP7 degradation because only K48 ubiquitin linkage can be recognized and processed by 26S proteasome system.

### ATM-mediated threonine 440 (T440) phosphorylation controlled the ubiquitination of LARP7

BRCT domain interactions are enhanced by serine or threonine phosphorylation (Yu et al., 2003). The increased interaction between LARP7 and the BARD1 BRCT domain indicated LARP7 was possibly phosphorylated during the DDR (Fig. 2d). The kinases ATM, ATR and DNAPK are drivers of phosphorylation cascade in the DDR(Ciccia and Elledge, 2010). To test if ATM, ATR or DNAPK mediated LARP7 phosphorylation as reported previously (Matsuoka et al., 2007), we suppressed their kinase activity with specific inhibitors and found that the ATM inhibitor KU55933, but not the ATR or DNAPK inhibitor, almost completely attenuated LARP7 decay in the IR- or CDDP-induced DDR (Fig. 3a, sFig. 6a). Depletion of ATM with a specific siRNA yielded the same outcome (sFig. 6b). ATM-mediated LARP7 phosphorylation was further supported by the finding that IR-induced phosphorylation of LARP7 occurred within SQ/TQ motifs that was specifically recognized by ATM or ATR (Fig. 3b). There are three SQ or TQ motifs in LARP7; among them, the TQ motif at positions 440-441 is a potential ATM-recognizing sequence, as predicted by Scansite 3.0 (https://scansite4.mit.edu/4.0/). This site is conserved across multiple species (Fig. 3c) and shares homology with a mouse sequence shown to be phosphorylated by IR (Matsuoka et al., 2007). We generated a custom antibody with validated specificity for phosphorylated T440 (sFig. 6c). Using this antibody, we demonstrated that IR induced a marked phosphorylation at T440 (Fig. 3d), and which was blocked by ATM inhibitor (Fig. 3e). Moreover, T440 substitution with alanine (T440A) abolished this phosphorylation (sFig. 6d). Together, these results demonstrated that ATM activation during the DDR catalyzes T440 phosphorylation on LARP7.

**Figure 3.**
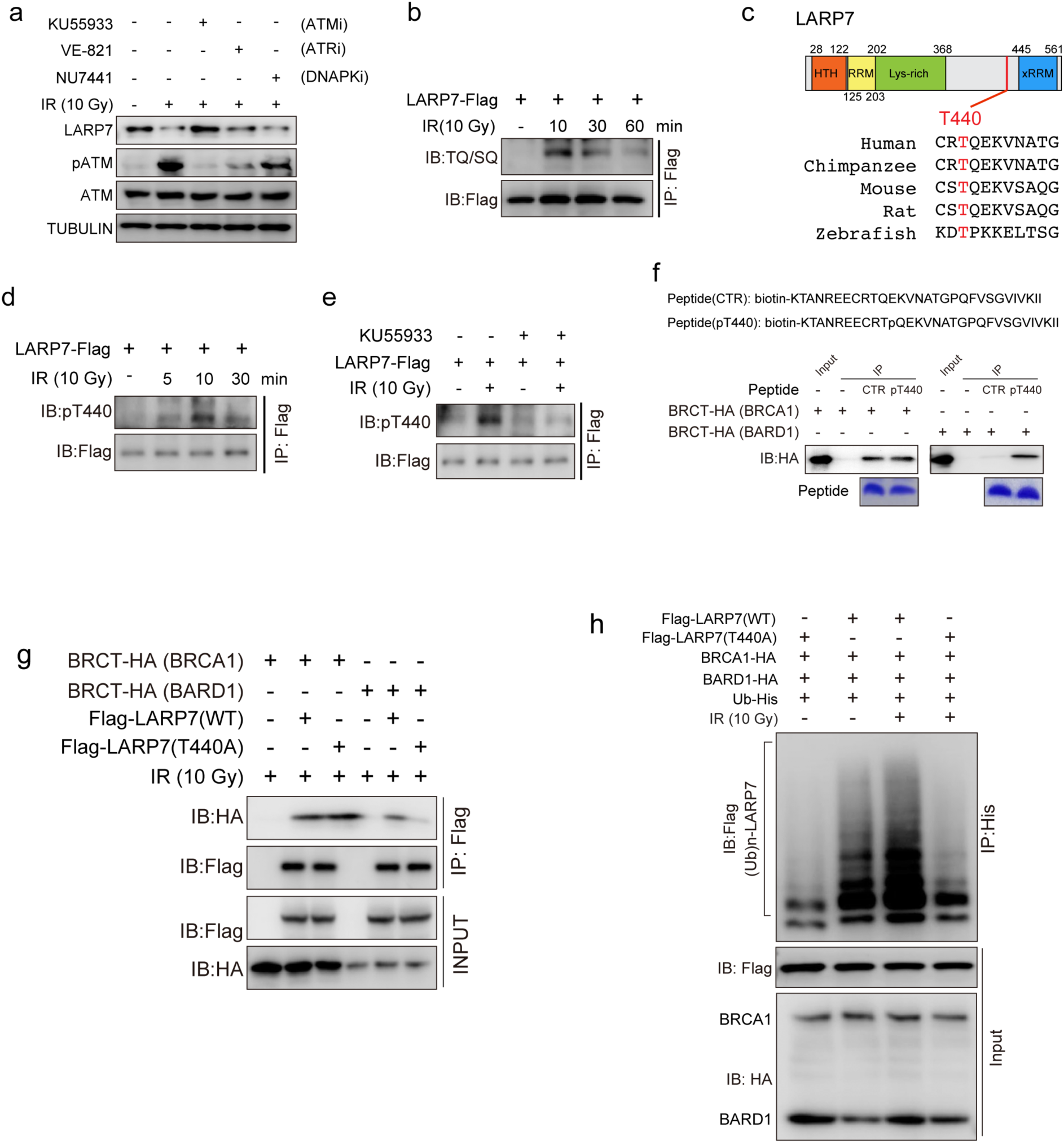
IR triggers ATM-mediated phosphorylation of LARP7 on T440. a, ATM phosphatase activity induced by DDR was required for LARP7 downregulation. KU55933: ATM inhibitor, 10 μM; VE-821: ATR inhibitor, 10 μM; NU7441: DNAPK inhibitor, 10 μM. b, IR induced phosphorylation at the ATM recognition motif of LARP7, as illustrated by a SQ/TQ phosphorylation antibody. c, T440 of LARP7 is evolutionarily conserved. d, T440 was phosphorylated after IR, as revealed by a phosphorylation-specific antibody. e, The ATM inhibitor attenuated IR-induced T440 phosphorylation. f, Immunoprecipitation assay showing that T440 phosphorylation enhanced the interaction of the LARP7 peptide with the BRCT domain of BARD1 but not with that of BRCA1. g, Co-IP results showing that T440A substitution attenuated the interaction of LARP7 with BARD1 but not with BRCA1. h, The T440A substitution attenuated IR-induced LARP7 polyubiquitination.

To evaluate if T440 phosphorylation could enhance the interaction of LARP7 with BRCA1/BARD1, we synthesized T440-phosphorylated and nonphosphorylated peptides and compared their affinities with BRCA1 or BARD1 BRCT domain using immunoprecipitation (Fig. 3e). The peptide with T440 phosphorylation had a much stronger interaction with BARD1 than the nonphosphorylated peptide. However, interactions of these two peptides with BRCA1 did not differ. Of note, the T440A replacement abrogated the IR-induced LARP7-BARD1 but not LARP7-BRCA1 interaction (Fig. 3g), and suppressed the polyubiquitination of LARP7 (Fig. 3h). Altogether, the results suggest that ATM-mediated T440 phosphorylation enhances LARP7-BARD1 interaction and facilitates BRCA1/BARD1-mediated LARP7 ubiquitination and degradation.

### LARP7 negatively regulated homology-directed repair (HDR)

DDR-mediated LARP7 degradation protected multiple cell lines including MDA-MB-231, a triple-negative breast cancer cell line, from DNA damage-induced cell death, which suggested LARP7 functioned as a negative DNA repair regulator to maintain the genome integrity (Fig. S7a). To test this hypothesis, we injured cells with IR and quantified chromosomal abnormalities including breaks, fusion, and ana- or polyploidy in wild-type or LARP7^-/-^ MEFs (Fig. 4a, sFig. 7b). There were significantly fewer chromosomal aberrations observed in LARP7^-/-^ cells than in wild-type cells, but the difference was abolished after recovery of LARP7 (Fig. 4a). Noteworthily, LARP7 restoration also resulted in more chromatid breaks and complex chromatid such as tri-radial chromosomes (Fig. S7b). IR-resulted DNA DSBs are predominantly repaired by either HDR or nonhomologous end joining (NHEJ) (Ciccia and Elledge, 2010). We assessed the effect of LARP7 on HDR using U2OS DR-GFP reporter cells, in which HDR reconstitutes the I-SceI restriction enzyme-disrupted GFP(Peng et al., 2009). The assay showed that LARP7 siRNA increased the GFP-positive cells (HDR repaired cells) by 2.07 fold comparing with control untreated cells, whereas LARP7 overexpression decreased it to 0.59 fold (p<0.05, Student’s t test, Fig. 4b-c). There results were reproduced in an analogous human embryonic stem cell based assay system (data not shown) (Xie et al., 2004). These data demonstrated that LARP7, as anticipated, negatively regulates HDR during DSB which is found to be substantially controlled by BRCA1 and BARD1.

**Figure 4.**
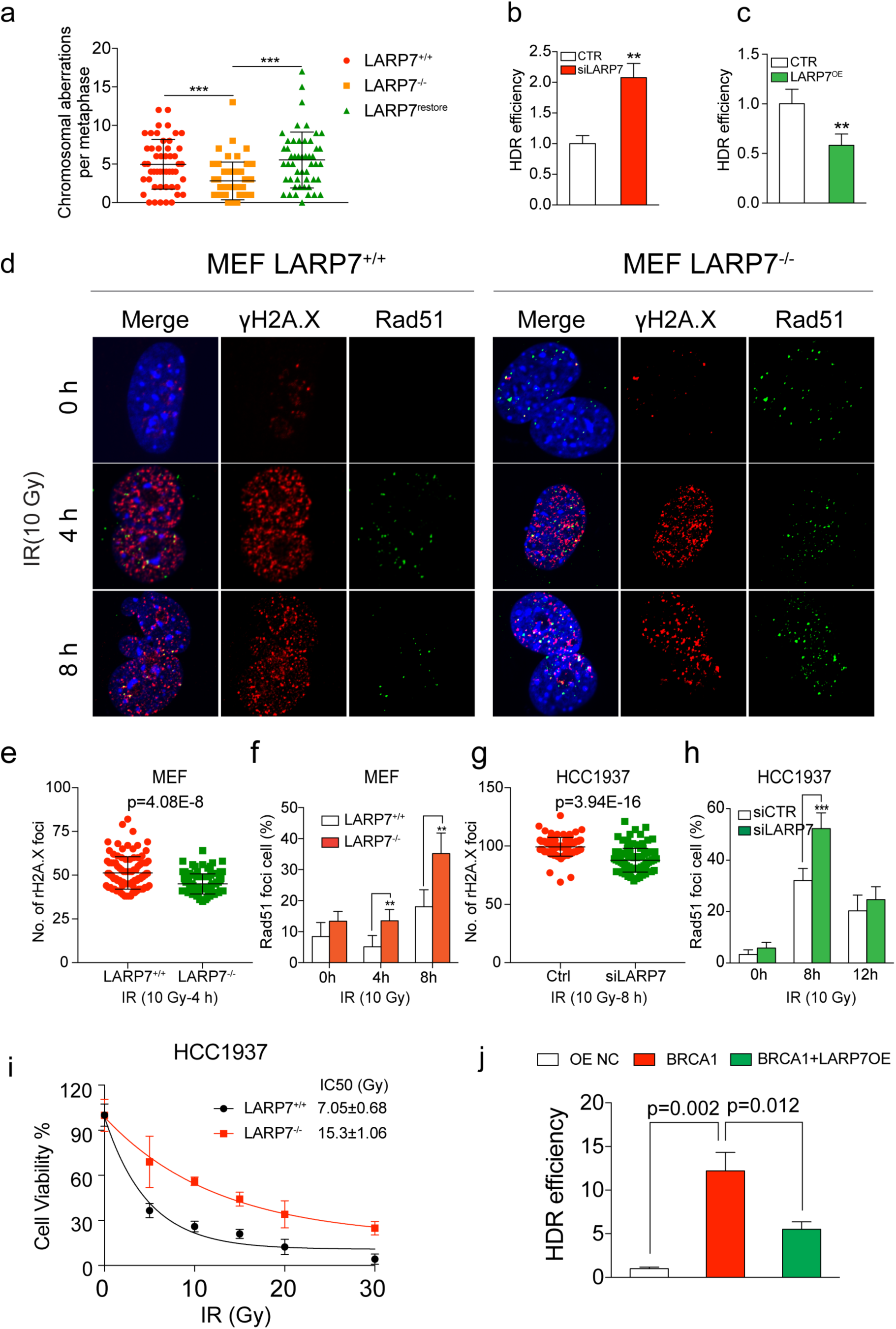
LARP7 impaired Rad51 recruitment and HDR. a, Karyotyping analysis of chromatin abnormalities in MEFs after IR (2 Gy). LARP7 knockout reduced the number of aberrant cells, while LARP7 restoration increased the number of cells with chromatin lesions. n=50 per group, Student’s t test, *: p<0.05, ***: p<0.001. b-c, HDR assay showing that knockdown of LARP7 increased the HDR capacity of U2OS DR-GFP reporter cells (b); conversely, overexpression of LARP7 impaired the HDR capacity (c). Each value is relative to the percentage of GFP+ I-SceI-transfected cells with control siRNA (b) or control vector (c) transfection, which was set to 1 and represents the mean ± SD of 3-5 independent experiments. Student’s t test, *: p<0.05, ***: p<0.001. d, Images showing Rad51 and γH2A.X foci in MEF nuclei after IR (10 Gy) at 0, 4 and 8 hours. LARP7 depletion enhanced Rad51 focus formation after IR in MEFs. γH2A.X and RAD51 double-positive foci indicate injured DNA regions under active repair. e, Statistical summary of the number of γH2A.X foci per cell in MEFs at 4 hours. Data: mean ± SD, Student’s t test, n=100 cells. f, Statistical analysis of the percentages of Rad51 focus-positive MEFs. Cells with more than 10 Rad51 foci were counted. Student’s t test, n=300-400, *: p<0.05, **: p<0.01. g, Statistical summary of the number of γH2A.X foci per cell in HCC1937 cells at 8 hours. Data: mean ± SD, Student’s t test, n=100 cells. H, Percentages of Rad51 focus-positive HCC1937 cells. Student’s t test, n=300-400, ***: p<0.01. i, CCK8 cell survival assay of wild-type and LARP7^-/-^ HCC1937. Data: mean ± SD, n=3. The IC50 was calculated with a one-phase decay model. j, HDR assay showing LARP7 attenuated BRCA1-mediated HDR. Mean ± SD, n=3-5, Student’s t test

γH2A.X and RAD51 form foci on chromatin lesions inside cells undergoing HDR (Ciccia and Elledge, 2010) (West, 2003). γH2A.X marks the DSB sites and recruits RAD51 to promote HDR (West, 2003). To gain further insight of the mechanism underlying LARP7-regulated HDR, γH2A.X and RAD51 focus formation upon IR was evaluated with immunofluorescence assays. IR significantly increased γH2A.X foci in wild-type MEFs, however, which was attenuated in LARP7^-/-^ MEFs (Fig. 4d-e). In consistence, the number of RAD51 focus-positive cells among LARP7^-/-^ MEFs was much higher at 4 and 8 hours than among wild-type MEFs (4 hours: p<0.01, 8 hours: p<0.05; Student’s t test, Fig. 4f). The increased RAD51 foci supported a conclusion that LARP7 depletion enhanced RAD51-meidiated HR and thereby reduced the DNA damage as indicated by γH2A.X foci.

The fact that LARP7’s degradation is mediated by BRCA1 and BARD1 suggested LARP7 worked at downstream effector of BRCA1/BARD1 to involve in HR process. To test it, we performed similar γH2A.X and RAD51 focus formation assay in HCC1937 breast cancer cell line that harbors a loss-of-function mutation in BRCA1 BRCT domain and found similar results to MEF that inhibiting LARP7 significantly increased RAD51 and decreased γH2A.X focus formation at 8 hours after IR (RAD51: p<0.001, rH2A.X: 3.94E-16, Student’s t test, Fig. 4g-h, sFig. 7c). LARP7 suppression also significantly increased the cell survival of HCC1937 cells treated with IR (Fig. 4i). Conversely, forced expression of LARP7 in MDA-MB-231 breast cancer cells harboring wildtype BRCA1 and low LARP7 suppressed RAD51 focus formation across all time points and increased γH2A.X foci (Fig. S7d-f), and meanwhile rendered more cells to death illustrated by cell survival assays and clonogenic assays (Fig. S8a-c). The independence of LARP7 on BRCA1 in regulating DNA damage demonstrated that LARP7 work downstream of BRCA1 to attenuate RAD51 assembly at DSB and negatively regulate HDR. By using DR-GFP HDR assays, we further illustrated LARP7 actually involved in a part of BRCA1-executed HDR by showing that overexpression of LARP7 in DR-GFP reporter cells compromised but not completely abolished the BRCA1’s capacity to promote HDR (Fig. 4j).

### LARP7 depletion deregulated CDK1 complex and induced G2/M cell cycle arrest

BRCA1 facilitates RAD51 assembly at chromatin damage sites and promoted HR {Zhao, 2017 #408} {Sy, 2009 #463}. Since our above experiments demonstrated LARP7 interacts with BRCA1/BARD1 complex, we first tested if BRCA1 mediated the enhanced RAD51 foci formation in LARP7 deletion cells. Treating MDA-MB-231 cells with IR for 6 and 12 hours induced significant BRCA1 foci at chromatin damage sites (sFig. 9a&b). But unlike RAD51, LARP7 deletion didn’t enhance BRCA1 focus formation at both tested time points. These results indicated LARP7 regulated RAD51 chromatin assembly possibly through a mechanism not depending on BRCA1. To further dissect out how LARP7 regulates RAD51 and HDR, we profiled the transcriptomes of three types of cells with or without LARP7 depletion by RNA sequencing. Genes regulating cell cycle especially G2/M checkpoints were significantly affected by LARP7 knockdown(sFig. 10a and Fig. 5a). Among them, CCNB1, CCNB2 and CDK1 (together: CCC), members of the CDK1 complex, were consistently downregulated in all LARP7-depleted cells. This downregulation was further confirmed by RT-PCR and WB analysis (Fig. 5b, sFig. 10b-g). Of note, depletion of LARP7 with siRNA or CRISPR suppressed the 7SK RNA which indicated a collapsed 7SK snRNP same to in IR- and CDDP-treated cells (Fig. S1a&d). Moreover, ectopic expression of LARP7 elevated CCC expression in MDA-MB-231 cells, suggesting that CCC are direct regulatory targets of LARP7 (sFig. 10h).

**Figure 5.**
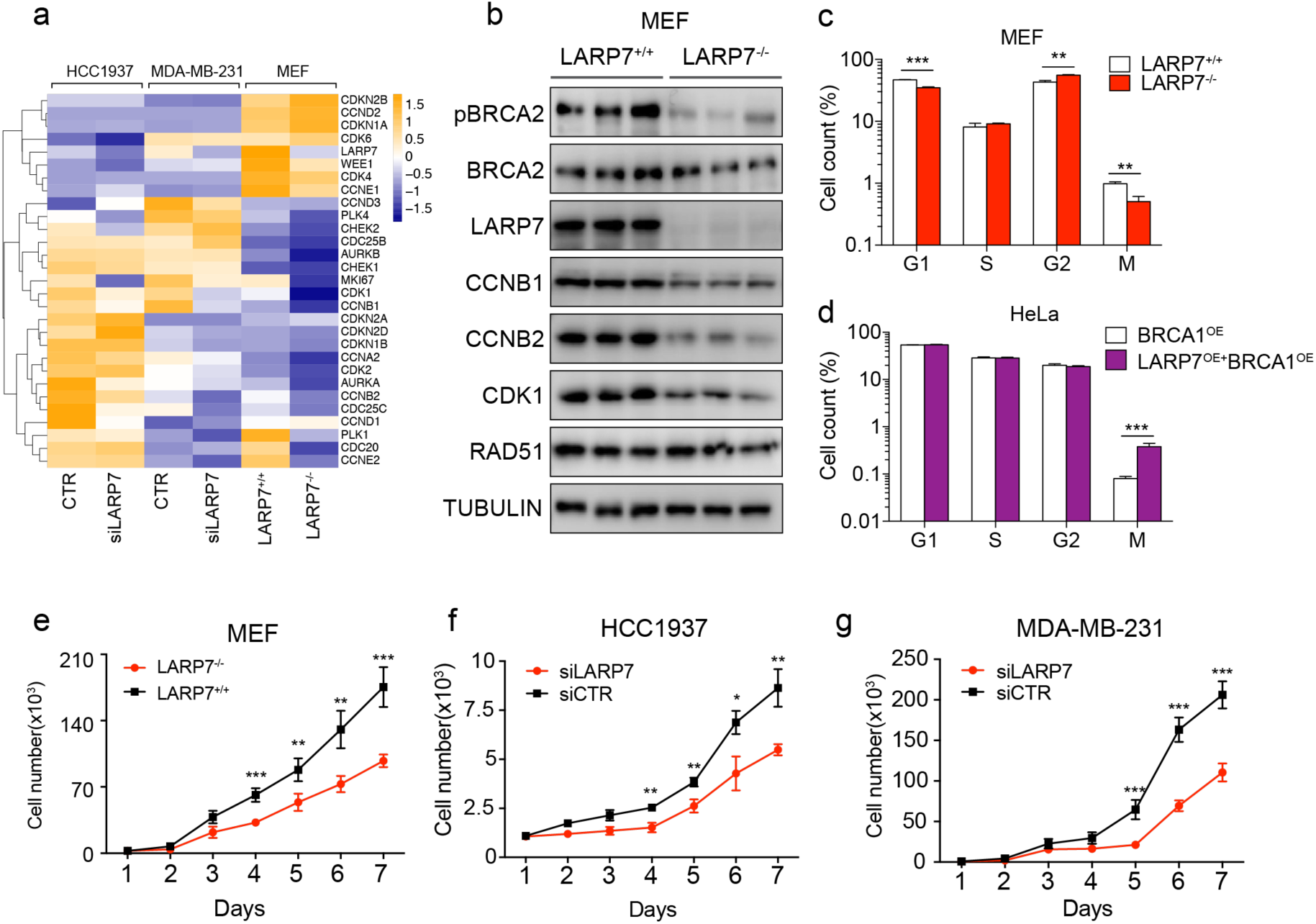
LARP7 depletion arrested cell at the G2/M transition. a, Expression profile of cell cycle genes in control and LARP7-depleted MEFs, HCC1937 cells and MDA-MB-231 cells as detected by RNA-seq. Reads per kilobase million (RPKM) values were normalized with Z-scores. b, LARP7^-/-^ MEFs had reduced protein levels of CCNB1, CCNB2 and CDK1, which are positive regulators of the G2/M transition, and reduced phosphorylation levels of BRCA2. c-d, FACS results of cell cycle distribution analysis with PI and PH3 double staining. LARP7 depletion induced cell cycle arrest at the G2/M transition in MEFs (c), while overexpression of LARP7 overcame the BRCA1-induced G2/M arrest (d). Data: mean ± SD, two-tailed student’s t test, n=3, **: p<0.01, **: p<0.01, ***: p<0.001. e-g, LARP7 depletion in MEFs (e), HCC1937 cells (f) and MDA-MB-231 cells (g) suppressed cell proliferation. Data: mean ± SD, Student’s t test, n=3-5, *: p<0.05, **: p<0.01, ***: p<0.001.

CCNB1 or CCNB2 together CDK1 promotes the G2/M transition during cell cycle progression (Pines and Hunter, 1989). The downregulation of CCC in LARP7-depleted cells suggested a defective G2/M transition and arrested cell cycle. We measured the cell cycle progression by dual-labeling four types of primary or cancer cells with propidium iodide and PH3, a specific indicator of M phase. LARP7-knockdown and LARP7-knockout cells had significantly lower proportions of cells in M phase and higher proportions in G2 phase in three tested cells lines except MDA-MB-231 cells, which indicated that these cells undergo the cycle arrest at the G2/M checkpoint (Fig. 5c, sFig. 11a-d). In contrast, the proportion of cells in S phase was not decreased, suggesting the G1/S transition, another other key cell cycle checkpoint, was largely unaffected by LARP7. BRCA1 blocks G2/M transition with mechanisms unclearly defined(Yarden et al., 2002). Ectopic expression of LARP7 overcame BRCA1-induced G2/M arrest in HeLa cells, indicating that BRCA1 induces the cell cycle arrest, if not all at least partially through it-mediated LARP7 degradation (Fig. 5d, sFig. 11e). Importantly, the LARP7 depletion-induced cell cycle also suppressed the growth of MEFs and other two breast cancer cell lines further supported LARP7’ s essence in cell growth (Fig. 5e-g).

### LARP7 depletion enhanced the RAD51-BRCA2 interaction and HDR by suppressing CDK1 complex

Recruitment of RAD51 to DNA lesions requires the assistance of BRCA2 (Esashi et al., 2005). CDK1 mediates BRCA2 phosphorylation and attenuates its interaction with RAD51 (Esashi et al., 2005). Thus, we hypothesized that the increased RAD51 focus formation in LARP7^-/-^ cells might result from reduced BRCA2 phosphorylation due to CCC downregulation. We therefore examined the levels of total and phosphorylated BRCA2 and RAD51 in LARP7-depleted MEFs, 293T, MDA-MB-231 and HCC1937 cells and observed markedly reduced BRCA2 phosphorylation compared to control, while total BRCA2 levels were unchanged (Fig. 5b and 6a, sFig. 10f-g). A mild decrease in RAD51 were also observed in HCC1937-knockout cells but not in MEFs, 293T or MDA-MB-231 cells (Fig. 5b and 6a, sFig. 10f-g). Conversely, upregulating LARP7 in MDA-MB-231 cells increased BRCA2 phosphorylation (sFig. 10h). Moreover, forced expression of the CDK1 complex in LARP7^-/-^ 293T cells increased BRCA2 phosphorylation, illustrating that attenuated expression of the CDK1 complex in fact accounted for the absent BRCA2 phosphorylation in LARP7-depleted cells (Fig. 6a). Decreased BRCA2 phosphorylation enhanced it’s interaction with RAD51, but was abrogated by restoration of CCC (Fig. 6b). LARP7 depletion significantly increased the RAD51 foci formation from 6h to 24h in HeLa cells comparing to wildtype cells, but addition of CCC in LARP7^-/-^ cells markedly abolished the increasement at all time points(Fig. 6c&d). Altogether, these data together with previous knowledge (Esashi et al., 2005) support a mechanism in which LARP7 disables RAD51 deposit at damage DNA and impairs HDR by upregulating a CCC-mediated BRCA2 phosphorylation.

**Fig. 6.**
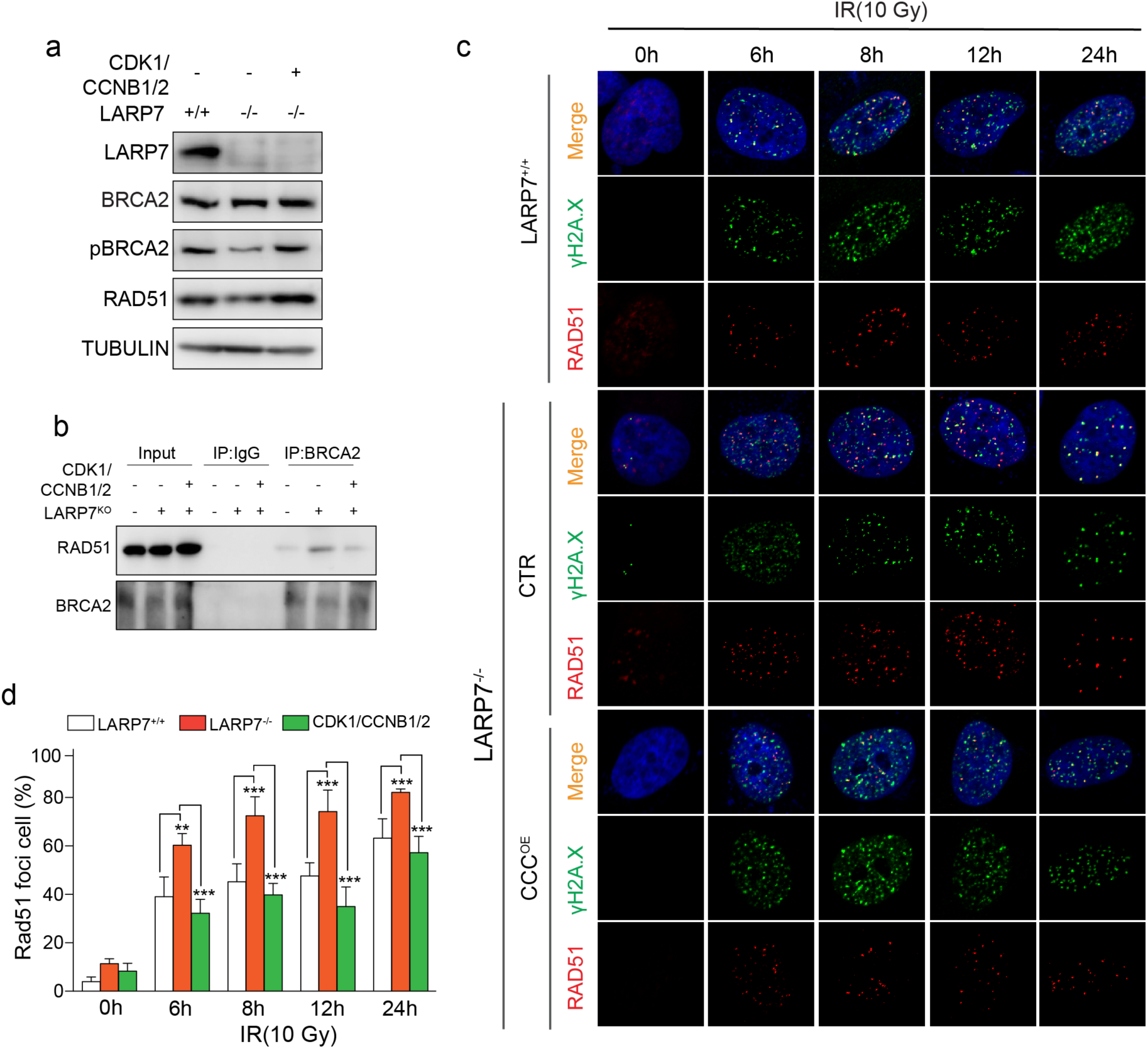
LARP7 depletion facilitated BRCA2-RAD51 interaction and RAD51 foci formation by downregulating CDK1 complex. a, LARP7 augmented the phosphorylation of BRCA2 in 293T cells, which was mediated by the CDK1 complex. b, The BRCA2-RAD51 interaction was enhanced in LARP7^-/-^ 293T cells, an effect that was reversed by restoration of the CDK1 complex. c, Immunofluorescence showing Rad51 focus formation upon IR treatment in LARP7^+/+^ HeLa cells, LARP7^-/-^ HeLa cells and LARP7^-/-^ HeLa cells with restored CCC. d, Statistical summary of Fig. 5C showing that the percentage of RAD51 focus-positive cells was higher among LARP7^-/-^ HeLa cells than among wild-type cells after IR. Addition of CCC abrogated this difference. Student’s t test, n≥300, *: p<0.05, **: p<0.01, ***: p<0.001.

### LARP7 promoted *in vivo* breast cancer tumorigenesis

The effects of LARP7 in promoting *in vitro* proliferation of breast cancer cells encouraged us to determine whether LARP7 regulates breast cancer tumorigenesis *in vivo*. Although LARP7 mRNA expression in breast cancer tissue is not different from that in normal tissue, as illustrated by the Gene Expression Profiling Interactive Analysis (GEPIA) database (Tang et al., 2017) (sFig. 12a), the levels of LARP7 protein were significantly higher in cancer tissue than in peritumoral normal tissue, as shown by immunochemical analysis of a total of 210 breast cancer tissue samples from patients (Fig. 7a, sFig. 12b and Table 4). Highly malignant tissues with TNM stages III to IV expressed more LARP7 than less-malignant (stage I) tissues. The tissue array data do not contain information regarding mutations in BRCA1, but BRCA1 expression was inversely correlated with LARP7 expression, which is consistent with the results obtained with the cell lines (Fisher’s exact test, p=0.008, Fig. 7b, sFig. 12c). We further investigated the role of LARP7 in tumorigenesis using the MDA-MB-231 cell line, a triple-negative breast cancer cell line with wild-type BRCA1, to generate xenograft tumors. Consistent with the *in vitro* growth results, the two LARP7^-/-^ cell lines created with two independent CRISPR small guide RNAs (sgRNAs) grew much slower in athymic nude mice than wild-type cells (p<0.001 from day 28 to 46, Student’s t test, Fig. 7c-d and sFig. 13a). To further examine if the effect of LARP7 on tumorigenesis was independent from BRCA1, as indicated *in vitro*, we compared the growth of BRCA1 mutant HCC1937 cell lines with or without LARP7 expression and found even greater growth inhibition in HCC1937 LARP7^-/-^ tumors (Fig. 7e and sFig. 13b-c). Proliferation markers including CCC were also significantly reduced in LARP7^-/-^ tumors, suggesting that LARP7 promotes tumorigenesis by regulating CDK1 complex expression (Fig. 7f and sFig. 13d). Accordingly, phosphorylation of BRCA2 was dramatically reduced in the LARP7^-/-^ xenografts, consistent with the cell line results (sFig. 13d). Collectively, these results demonstrate that high expression of LARP7 in breast cancer contributes to *in vivo* tumor growth and progression.

**Figure 7.**
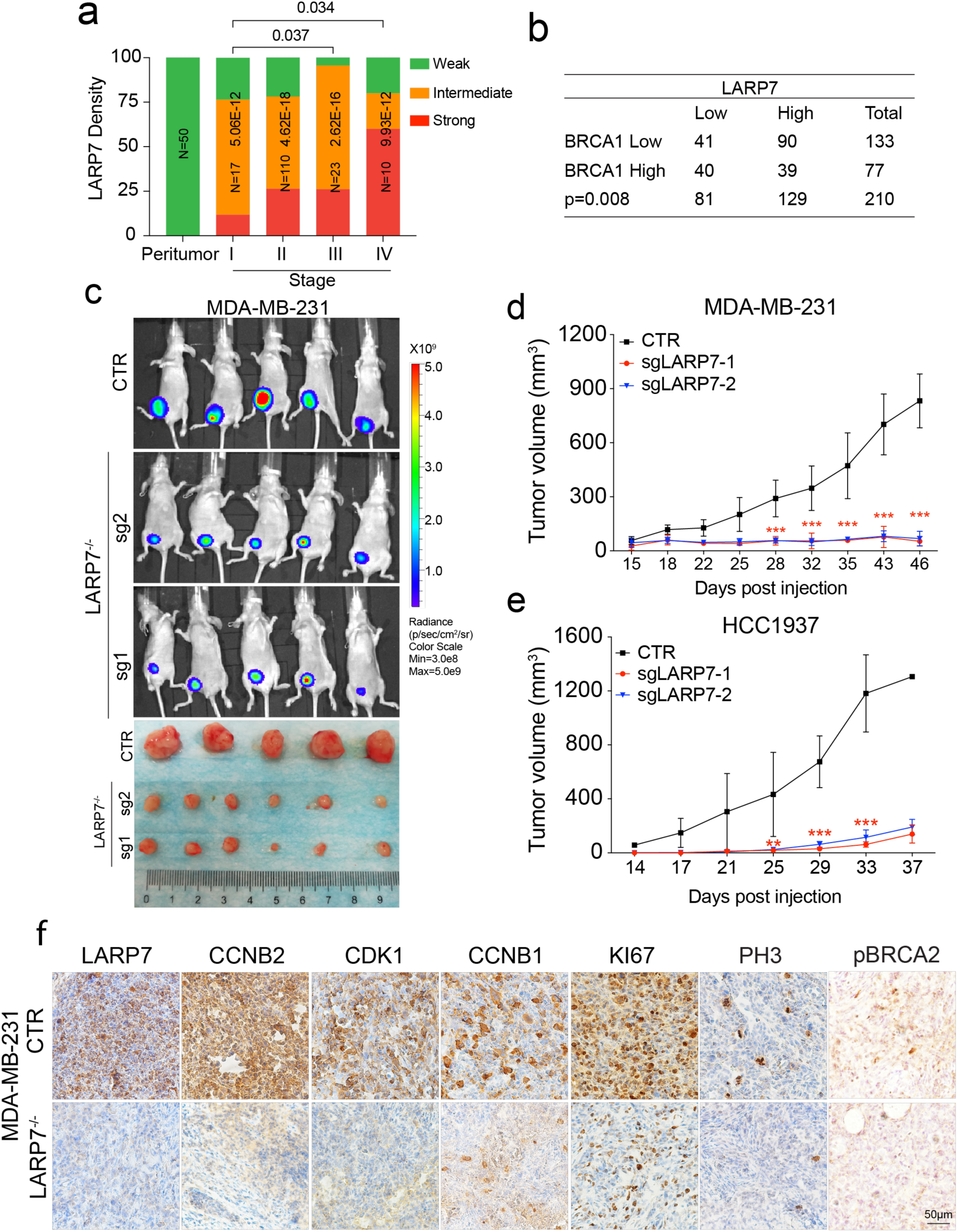
LARP7 promoted breast cancer tumorigenesis. a, LARP7 protein expression was elevated in breast cancer tissue, especially at late TNM stages, as revealed by LARP7 IHC. Wilcoxon rank-sum test, n=210; p<0.05 indicated significance. b, LARP7 levels were negatively correlated with BRCA1 protein levels in 210 breast cancer samples. Chi-squared test; p<0.05 indicated significance. c, LARP7 knockout suppressed xenograft tumor growth from triple-negative cancer (TNC) MDA-MB-231 cells. The top panel shows luminescence images of transplanted tumors. The bottom panel shows images of harvested tumors. Sg1 and sg2 indicate the LARP7^-/-^ MDA-MB-231 tumors generated with two independent sgRNAs. d, Growth curve of wild-type and LARP7^-/-^ MDA-MB-231 xenograft tumors. Mean±SD, Student’s t test, n=5 mice for wild-type, n=6 mice for LARP7^-/-,^ *: p<0.05, **: p<0.01. e. Tumor growth curve of HCC1937 xenografts. Mean±SD, Student’s t test, n=6 mice for CTR, n=7 for the LARP7^-/-^ groups, *: p<0.05, **: p<0.01, ***: p<0.001. f. IHC images of wild-type and LARP7^-/-^ MDA-MB-231 xenografts. pBRCA2: phosphorated BRCA2 (Ser3291).

### LARP7 depletion increased the therapy resistance and was related to the relapse of breast cancer

Chemoradiotherapy resistance is a primary cause of tumor recurrence and lethality in the clinic. The IR and CDDP resistance shown *in vitro* in LARP7-depleted cell lines compelled us to examine if LARP7 contributes to the *in vivo* therapy resistance. In line with this hypothesis, four cohort studies illustrated that recurrent breast cancers had lower LARP7 levels than non-recurrent breast cancers (sFig. 14a). In addition, breast cancer patients with low LARP7 levels had much shorter relapse-free survival (RFS) times after chemotherapy than patients with high LARP7 levels (average survival in months: 34 vs. 77.28, p=0.022, logrank test; data from KM Plotter, Fig. 8a) (Lanczky et al., 2016). The average survival time (months) of LARP7-low patients was comparable to that of BRCA1-high patients and was even shorter than that of BRCA2-high patients (sFig. 14b-c). BRCA1 and BRCA2 are canonical genes leading to chemotherapy resistance, so the comparable or even worse RFS in LARP7-patients suggested a critical role of LARP7 in regulating therapy resistance. Noteworthily, no difference was observed in RFS between LARP7-high and LARP7-low patients that both had received endocrine therapy (sFig. 14d), suggesting that LARP7 regulates therapy resistance through a specific mechanism of DNA damage repair.

**Figure 8.**
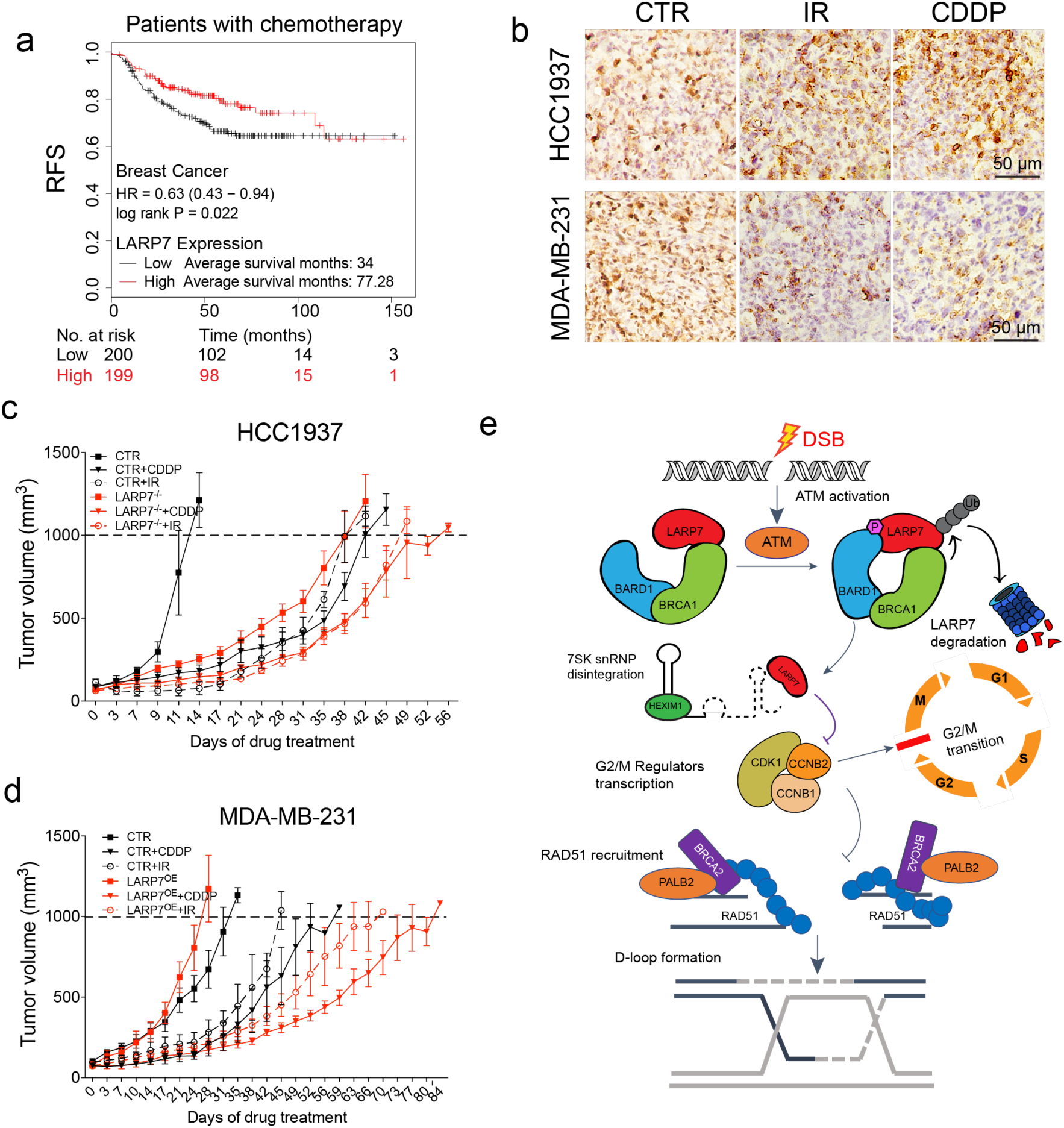
LARP7 enhanced genotoxic therapy sensitivity. **a.** Breast cancer patients receiving chemotherapy were stratified by LARP7 expression (LARP7 High= top 25%; LARP7 Low=bottom 25%), and RFS was analyzed by the Kaplan-Meier method. HR: hazard ratio. The numbers of patients that died at different times are labeled at the bottom of the plots. Statistical significance was calculated with logrank tests, and p<0.05 indicated significance. b. IHC showing the LARP7 protein levels in CDDP- and IR-treated tumors. CDDP and IR induced significant downregulation and extranuclear shuttling of LARP7 in MDA-MB-231 xenograft tissue but not in HCC1937 xenograft tissue. c. Tumor sizes of wild-type or LARP7^-/-^ HCC1937 xenografts treated with CDDP or X-rays. Wild-type tumors with or without treatment: n=7, LARP7^-/-^ tumors with or without treatment: n=8, mean±SD. The intersection of each curve with the dotted line indicates the time (from treatment initiation) at which the tumor size reached 1000 mm^3^. d. Tumor sizes of wild-type or LARP7^OE^ MDA-MB-231 xenografts treated with CDDP or X-rays. Wild-type tumors with or without treatment: n=7, LARP7^OE^ tumors with or without treatment: n=7, mean±SD. The intersection of each curve with the dotted line indicates the time (from treatment initiation) at which the tumor size reached 1000 mm^3^. e. Schematic of the mechanism by which LARP7 attenuated HDR. DNA DSBs activated ATM, which phosphorylated LARP7 at T440, increased the interaction of LARP7 with BARD1 and triggering the ubiquitination and degradation of LARP7 through the 26S proteasome. Under ordinary conditions, LARP7 promotes the expression of CCC. Depletion of LARP7 suppressed CCC expression, which arrested the cell cycle at the G2/M checkpoint. In addition, LARP7 depletion decreased BRCA2 phosphorylation, which enhanced RAD51 recruitment to DNA damage sites and thus increased RAD51-mediated homologous recombination repair.

To further validate the role of LARP7 in chemotherapy resistance, we first established xenograft tumor models using HCC1937 cells, which are chemoradiotherapy sensitive due to a loss-of-function mutation in BRCA1, and treated them with CDDP and IR. Immunochemistry showed that LARP7 levels were not altered by these treatments which was similar to the results *in vitro* (Fig. 8b). However, LARP7 depletion significantly dampened the susceptibility of xenografts to CDDP, as revealed by attenuated increases in time-to-endpoint (TEP: the time from treatment initiation to the point at which the tumor size reached 1000 mm^3^) in LARP7^-/-^ xenograft mice (TEP: average 39.6 days in wild-type mice vs. 17.6 days in LARP7^-/-^ mice, p<0.001, unpaired t test, Fig. 8c and sFig. 14e-f). Similar resistance was also observed in IR treatment which is mirror to clinical radiotherapy (TEP: average 38.9 days in wild-type mice vs. 21.6 days in LARP7^-/-^ mice, Fig. 8c and sFig. 14e-f).

MDA-MB-231, a triple-negative breast cancer cell line, is resistant to hormone therapy and ample types of chemotherapies. MDA-MB-231 cells express intact BRCA1/BARD1 and low levels of LARP7. To determine whether MDA-MB-231 cells could regain therapy sensitivity after upregulation of LARP7, as shown in the *in vitro* assays, we compared the tumor growths in MDA-MB-231 xenografts with or without expression of LARP7 after treatment with CDDP and IR. Both CDDP and IR induced marked reductions and extranuclear shuttling of LARP7 in MDA-MB-231 cells (Fig. 8b). Ectopic expression of LARP7 promoted *in vivo* tumor growth (Fig. 8d and sFig 14g). However, after treatment with either CDDP or IR, LARP7^OE^ tumors grew even more slowly than control tumors (Fig. 8d and sFig. 13e). The increase in TEP due to treatment was significantly greater in LARP7^OE^ tumors than in wild-type tumors (wild-type vs. LARP7^OE^: 31.7 vs. 71.3 days for CDDP, 27.8 vs. 50.9 days for IR; sFig. 14h). LARP7^OE^ tumors also had more DNA damage gauged by γ-H2AX and apoptosis gauged by active caspase-3 staining (sFig. 14i). These results together reflected a better susceptibility of LARP7^OE^ MDA-MB-231 tumors to CDDP or IR.

## Discussion

We report BRCA1/BARD1-regulated mechanism centered on LARP7 that coordinates G2/M arrest with HDR-mediated DNA repair. ATM activation by DSBs induces phosphorylation of LARP7 at T440, increasing LARP7 binding to and K48 polyubiquitination by BRCA1/BARD1. This sends LARP7 to the 26S proteasome for degradation and leads to a collapse of 7SK snRNP. 7SK snRNP disassembly suppresses the expression of the CDK1 complex, which arrests the cell cycle at the G2/M checkpoint. Suppression of the CDK1 complex also reduces BRCA2 phosphorylation, which enhances RAD51 recruitment to damaged DNA and promotes homologous recombination-directed DNA repair (Fig. 8e). This is a new pathway for BRCA1/BARD1-mediated HR which is centered on 7SK snRNP and has not been reported before.

7SK snRNP is the hub of RNAPII pausing regulatory network. It recruits HEXIM1, 2 to jail CDK9, the catalytic unit of pTEFb, prevents it to catalyze the serine 2 phosphorylation on RNAPII CTD domain and thus pauses the engaged RNAPII at the TSS proximal regions (Krueger et al., 2008) (Nguyen et al., 2001) (Yang et al., 2001). To release RNAPII pausing and activate the transcription, it acquires Bromodomain-containing protein 4 (BRD4) (Bisgrove et al., 2007)(Yang et al., 2005), HIV associated Tat1 protein (Barboric et al., 2007) or other regulators such as acetylated ETS1 which was recently defined by our group (Chen et al., 2017), to release the pTEFb from 7SK snRNP and relocate it to the chromatin. It has been recently recognized that there is a crossroad between two pathways of RNAPII pausing and DNA repair. A RECQ family of helicase of RECQL5 repress RNAPII elongation and prevents the genome instability caused by transcription stress (Saponaro et al., 2014). The inhibitors of BRD4 and CDK9, two key components constituting the activate pTEFb, suppress the expression of C-terminal binding protein (CtBP) interacting protein (CtIP), impairs DNA end resection and HDR (Sun et al., 2018). Their combination with PARP inhibitors exhibited the synergistic therapeutic effect and reversed the resistance of multiple type of tumors to PARP inhibitors.

7SK snRNP, although not directly associated with RNAPII, was also found to involve in DDR. A very recent study reported that the genotoxic chemical 4-nitroquinoline 1-oxide (4-NQO), a mimetic of UV inducing the nucleotide excision repair (NER), increased RNA-binding motif protein7 (RBM7) bind to 7SK snRNP, which in turn released pTEFb and promoted RNAPII pause release(Bugai et al., 2019). Likewise, our study demonstrate LARP7, the core component of 7SK snRNP response to IR- and CDDP-induced DDR and attenuates HDR. Although both studies unveiled the importance of 7SK snRNP in the response to variant types of genotoxic stress, the mechanism reports seems different. We revealed the underlying mechanism that IR and CDDP induced LARP7 degradation and the disruption of 7SK snRNP, which silenced the cell cycle and improved the HDR. In contrast, Andrii Bugai’s study suggested 4-NQO induced RBM7’s relocation to 7SK snRNP to releases pTEFb and promoted the cell survival, while the integration of 7SK snRNP remains unaffected throughout the response process (Bugai et al., 2019). We reasoned the difference is possibly because of the distinct genotoxic stressed applied in two studies. IR and CDDP is reported to predominantly induce the ATM-mediated DDR, while NER activates the ATR pathway (Ciccia and Elledge, 2010), and our results illustrated that BRCA1-mediated LARP7 ubiquitination and degradation depends on ATM rather ATR. Thus, the activation of different DDR pathways might account for the discriminative discovery between two studies, but whether this is the case requires further validation. Nevertheless, both studies convergently underpinned the importance of RNAPII pausing pathway activation mediated by 7SK snRNP in the cell’s response to DNA damage.

Uncontrolled cell cycle progression is the hallmark of tumor cells. HDR predominantly occurs shortly after DNA replication in the G2 phase, when sister chromatids are more available. Equal to its key role in HDR, BRCA1 also controls the G2/M checkpoint to avert the aberrant chromosome segregation (Huen et al., 2010). Depletion of RAP80 or ABRAXAS, which recruit BRCA1 to DNA damage foci, impaired BRCA1-dependent G2/M arrest, but the phenotype and magnitude of cell growth arrest in these cells was mild in comparison to BRCA1-deficient cells (Sobhian et al., 2007) (Kim et al., 2007). During DDR, BRCA1 also regulates the phosphorylation status and activation of CHK1 to control the G2/M transition (Yarden et al., 2002). However, mechanism by which the ubiquitinase activity of BRCA1/BARD1 regulates cell cycle arrest was previously unknown. LARP7 had been shown to regulate cell growth, and homozygous LARP7 loss-of-function results in the primordial dwarfism that is mainly caused by cell growth arrest (Alazami et al., 2012). Here, we show that LARP7 enhanced expression of key CDK1 subunits and activated the G2/M checkpoint. BRCA1-mediated K48 ubiquitination and depletion of LARP7 suppressed cell growth *in vitro* and xenograft tumor progression *in vivo*. These lines of evidence together show that BRCA1 ubiquitinase activity is integral to BRCA1-mediated G2/M checkpoint control, and that LARP7 is a key BRCA1 ubiquitination substrate and downstream effector.

In summary, this study identifies LARP7 as a novel oncogene that can double gate cell cycle progression and genome stability of tumor cells. Targeting LARP7 and 7SK snRNP pathway could become a promising path to prevent tumor or increase the therapeutic efficacy as manifested by our *in vitro* and *in vivo* results.

## STAR★ METHODS

Detailed methods are provided in the online version of this paper and include the following:

- KEY RESOURCES TABLE
- CONTACT FOR REAGENT AND RESOURCE SHARING
- EXPERIMENTAL MODELS AND SUBJECT DETAILS

- Experimental animals
- Xenograft model
- Chemo- and radiotherapy
- METHOD DETAILS

- Cell culture
- Virus production and infection
- Cell viability assay
- Cell proliferation assay
- Colony formation assay
- Protein purification and GST pulldown assay
- Western blotting
- In vitro ubiquitination assay
- In vivo ubiquitination assay
- Bimolecular fluorescence complementation (BiFC) assay
- Coimmunoprecipitation
- Protein half-life assay
- Immunohistochemistry
- Immunofluorescence
- RNA-seq
- Cell cycle analysis
- Chromosomal abnormalities
- Homologous repair assay
- Computational analysis

## SUPPLEMENTAL INFORMATION

Supplemental information includes eight figures and four tables.

## AUTHOR CONTRIBUTIONS

B.Z. conceived this study. F.Z., P.Y.Y., H.Y.L., H.Y.L, Z.X.L., S.Y.Z., F.Z., H.J.Y., X.D.L., L.L. and

W.W.T performed the bench experiments. J.H.C. performed the RNA-seq analysis. X.J. assisted with the karyotyping experiment. A.Y.X. provided ESC DR-GFP cells and assisted with the HDR experiment. W.T.P, K.S., Z.G.H, S.C, Y.W.C.,B.X.G and B.Z. reviewed and revised the paper. F.Z., P.Y.Y., H.J.Y. and B.Z. wrote the paper.

## ACKNOLEDGEMENTS

We thank Dr. David Clapham and Dr. Sean Li (Boston Children’s Hospital) for providing the HeLa and various other cancer cell lines and Dr. Shiaw-Yih Lin for providing the U2OS DR-GFP reporter cell lines. B.Z. is funded by the National Science Foundation of China (91539109, 31671503 and 31872836), a Thousand Young Talents Award (16Z127060017), the Innovation Program of the Shanghai Municipal Education Commission (2017-01-07-00-01-E0028), and the National Key Research and Development Program of China (2018YFC1312702 and 2018YFC1002400).

## DATA AVAILABILITY

Raw and processed RNA-seq data were deposited to GSE124393

## COMPETING INTERESTS

The authors declare no competing interests

## Methods

### Animals

Female nude mice (CByJ.Cg-Foxn1nu/J, Jax #000711) 5-6 weeks of age were purchased from Lingchang BioTech for breast cancer xenograft studies. A LARP7^-/-^ conventional knockout mouse model was generated with CRISPR techniques in JAX mice (sFig. 1e-g) bred on a C57BL6 background. The mice were housed under standard conditions with a 12-hour light/dark cycle and access to food and water ad libitum. *In vivo* tumorigenesis, chemotherapy and IR treatment were performed in accordance with animal protocols approved by the Institutional Animal Care and Use Committee of Shanghai Jiao Tong University.

### Xenograft model

Exponentially growing breast cancer cells including MDA-MB-231-luc-LARP7^-/-^ (3×10^6^), MDA-MB-231-luc-sgRNA-control (CTR) (3×10^6^), HCC1937-luc-LARP7^-/-^ (5×10^6^) and HCC1937-luc-sgRNA-CTR (5×10^6^) cells were mixed with 50% Matrigel and injected subcutaneously above the right hind leg of 6- to 8-week-old female nude mice. Beginning two weeks post injection, tumor size was measured every 3-4 days. The tumor volume was measured in two dimensions, length and width, using electronic calipers and was calculated by the formula volume= (length X *width*^2^)/2. Once the tumor volumes were >1000 mm^3^, they were first assessed for luciferase activity by intraperitoneal injection of D-luciferin (150 mg/kg) and imaging with a Xenogen IVIS-2 Imaging System; then, the tumors were dissected out, weighed and imaged. Dissected cancer tissue was further subjected to paraffin embedding for immunohistochemistry (IHC) assays.

### Chemo- and radiotherapy

Exponentially growing cells including MDA-MB-231-luc-LARP7^OE^ (3×10^6^), MDA-MB-231-luc-CTR (3×10^6^), HCC1937-luc-LARP7^-/-^ (5×10^6^), and HCC1937-luc-sgRNA CTR (5×10^6^) cells were mixed with 50% Matrigel and injected subcutaneously above the right hind leg of the mice. Two weeks after injection, tumor volumes were externally measured in two dimensions using electronic calipers (volume=[length× width^’^]/2). When the tumor volumes reached approximately 100 mm^3^, the mice were treated with vehicle or CDDP (Sigma-Aldrich) intraperitoneally at doses of 5 mg/kg/week or with radiation (5 Gy for HCC1937 xenografts and 10 Gy for MDA-MB-231 xenografts) (X-ray Irradiator, Rad Source Technologies, RS2000). The animals were sacrificed by CO_2_ asphyxiation when the tumor volumes were >1000 mm^3^ or if the animals were obviously ill. Then, the tumors were dissected out and subjected to paraffin embedding for IHC assays.

### Cells

HEK293T cells were provided by William T. Pu (Harvard University), HeLa cells were provided by David E. Clapham (Harvard University), and MDA-MB-231 and MCF7 cells were provided by Dr. Sean Li (Harvard University). HCC1937, MDA-MB-436, MDA-MB-468 and HBL-100 cells were purchased from the Cell Bank of the Shanghai Institutes for Biological Sciences. MEFs were purified from LARP7^+/+^ or LARP7^-/-^ embryos at the age of E13.5 and were used for experiments between passages 1 and 6. All cell lines were confirmed to be free of mycoplasma contamination.

HBL-100 cells were grown in RPMI 1640 medium (Gibco) supplemented with 10% fetal bovine serum (FBS). HEK293T and HeLa cells were grown in DMEM (Gibco) supplemented with 10% FBS. MCF7 cells were grown in MEM (Gibco) supplemented with 1.5 g/L NaHCO3, 0.11 g/L sodium pyruvate, mg/ml bovine insulin, and 10% FBS. HCC1937 cells were grown in RPMI 1640 medium (Gibco) supplemented with 1.5 g/L NaHCO3, 2.5 g/L glucose, 0.11 g/L sodium pyruvate, and 10% FBS. MDA-MB-231 and MDA-MB-468 cells were grown in L-15 medium (Gibco). MDA-MB-436 cells were grown in L-15 medium supplemented with 10 μg/ml insulin, 16 μg/ml glutathione, and 10% FBS. U2OS DR-GFP reporter cells were grown in RPMI 1640 medium supplemented with 1.5 g/L NaHCO3, 2.5 g/L glucose, 0.11 g/L sodium pyruvate, and 10% FBS. ESCs were cultured with irradiated MEF feeders in DMEM supplemented with 10% FBS and 1000 µg/ml LIF. All of the above cells were maintained at 37°C under 5% CO_2_ in a humidified incubator with the exception of MDA-MB-231, MDA-MB-436, and MDA-MB-468 cells, which were cultured in 100% regular air.

### Stable cell lines

To generate stable luciferase reporter cell lines, MDA-MB-231 and HCC1937 cells were infected with lentivirus packaged in the pLenti-CMV-v5-luc-Blast plasmid, and monoclonal cells were picked after selection with blasticidin (ant-bl-05, InvivoGen). The resulting reporter cell clones (MDA-MB-231-luc and HCC1937-luc) were validated by measuring the luciferase activity after adding D-luciferin (Promega) to the cell culture medium. To further generate cell lines stably over- or underexpressing LARP7, MDA-MB-231 and HCC1937 cells were infected with lentivirus with a Lenti-LARP7-Flag or pLKO.1-shLARP7 vector, respectively, for two days and then cultured with puromycin (2 μg/ml for MDA-MB-231 cells and 1 μg/ml for HCC1937 cells) for 7 days. Single clones were then picked to 96-well plates, and the concentration of puromycin was changed to 0.5 μg/ml. The surviving stable cells were maintained and propagated in culture medium supplemented with puromycin (1 μg/ml). Puromycin was withdrawn 2 passages prior to mouse injection to eliminate adverse effects of residual puromycin.

To generate LARP7 homozygous knockout cell lines, 293T, HeLa, HCC1937-luc and MDA-MB-231-luc cells were infected with lentiviruses with the LentiCRISPR v2-sgLARP7 vector (sequences designed at http://crispr.mit.edu, Table 2) for 48 hours and selected with puromycin (2 μg/ml for MDA-MB-231 cells and 1 μg/ml for HCC1937 cells) for 7 days. Monoclones were picked for further propagation. Homozygous knockout was confirmed by PCR and WB analysis.

### Virus production and infection

For overexpression of LARP7, LARP7 fused with Flag was cloned into a pLenti-CMV-puro vector (17452, Addgene) to generate Lenti-LARP7^OE^ vectors. As a control, GFP was cloned into the same vector. To knock out LARP7, a 21 nt sgRNA was cloned into a LentiCRISPR v2 vector (52961, Addgene) to generate LentiCRISPR v2-sgLARP7. The purified target vectors together with the packaging vector pMD2.G and psPAX2 (Addgene, 12259 and 12260) were transfected into 293T cells using PEI (polyethylenimine, biopolymer). The cell culture medium containing virus particles was collected on days 2 and 3 after transfection and purified with 9% PEG6000 through centrifugation. The virus titer was estimated with a Lenti-X™ p24 Rapid Titer kit by measuring the viral envelope protein p24. A total of 1-5 IU (infection units) of lentivirus were added to the cell culture medium for infection supplemented with 10 μg/ml of polybrene (Sigma-Aldrich). The infection medium was replace with fresh medium 4-8 hours after infection.

### Cell viability assay

LARP7-knockout, LARP7-knockdown, LARP7-overexpressing or control cells (5000 cells for CDDP; 2000 cells for IR) were placed in 96-well plates; 24 hours after seeding, the cells were treated with the indicated doses of IR (X-ray Irradiator, Rad Source Technologies, RS2000) or CDDP (Sigma-Aldrich, P4394). An Enhanced Cell Counting Kit-8 (CCK8; Beyotime, C0042) was used to measure cell viability according to the manufacturer’s instructions. Briefly, 10 μl of CCK8 solution was added to each well. The plates were incubated at 37°C for 1 hour and then measured in a microplate reader at a wavelength of 450 nm.

### Cell proliferation assay

MEFs (2500 cells/well), MDA-MB-231 cells (1000 cells/well), and HCC1937 cells (1000 cells/well) were seeded into 96-well plates. At the indicated detection times, 10 μl of CCK8 reagent (Beyotime, C0042) was added into each well. The plates were incubated at 37°C for 1 hour, and then the optical density was detected at a wavelength of 450 nm.

### Colony formation assay

LARP7-knockout or LARP7-overexpressing MDA-MB-231 (1000 cells per well) and HeLa cells (250 cells per well) were plated to six-well plates, irradiated with the indicated doses of X-rays or incubated with CDDP (Sigma-Aldrich, P4394) for 24 hours, and then further cultured for 7-10 days. After incubation with a fixing/staining solution (0.05% crystal violet, 1% formaldehyde, 1% methanol) for 20 min at room temperature, the colonies were imaged under a stereomicroscope and counted with ImageJ.

### Protein purification and GST pulldown assay

His-tagged LARP7 and GST-tagged BRCT domains of BRCA1 and BARD1 were transfected into BL21/DE3 *E. coli* grown in LB medium and induced by addition of 0.25 mM IPTG. To purify the BRCA1/BARD1 heterodimer RING domain, His-tagged N-terminals of BRCA1 (residues 1-304; Addgene, 12645) and BARD1 (residues 26-327; Addgene, 12646) were cotransfected into BL21/DE3 *E. coli*. The bacterial pellet was lysed with a Constant Cell Disruption System in buffer A (20 mM Tris-Cl pH 7.4, 150 mM NaCl, 2 mM MgCl2, and 5 mM imidazole) followed by supernatant clarification and His-tagged protein capture using TALON Metal Affinity Resin (Clontech, 635501). The TALON Metal Affinity Resin with captured proteins was washed with buffer B (20 mM Tris-Cl pH 7.4, 150 mM NaCl, and 10 mM imidazole) three times and then eluted with buffer C (20 mM Tris-Cl pH 7.4, 150 mM NaCl, 200 mM imidazole). A final change to imidazole-free buffer was performed with an Amicon column (10K or 30K, Millipore).

For the GST pulldown assay, 5 µg of purified LARP7 protein was added to glutathione Sepharose 4B beads bound to GST, GST-BRCA1-BRCT, or GST-BARD1-BRCT and incubated in binding buffer (1x PBS pH 7.4, 1% Triton X-100, 1 mM PMSF, 1 mM NaVO3, 1x cocktail, 1x NaF) at 4°C overnight. After 3 washes with binding buffer, the beads were boiled in 100 µl of 2x SDS-PAGE loading buffer. The samples were then analyzed via western blotting.

### Western blotting

Washed cell pellets were lysed in high-salt buffer B (20 mM HEPES pH 7.9, 450 mM NaCl, 25% glycerol, mM EDTA, 0.5 mM DTT, 1% NP40 and 0.5% SDS plus protease and phosphatase inhibitors) for 15 min at 4°C and centrifuged at 12000 rpm for 10 min. The supernatant was boiled with 4x Laemmli buffer. The extracted proteins were separated on SDS-PAGE gels, transferred to PVDF membranes and probed with primary antibodies (See Table 1) overnight at 4°C. The bound proteins were visualized with enhanced chemiluminescence substrate (Immobilon Western, WBKLS0500, Millipore) and imaged with an Amersham Imager 600.

To validate the specificity of the pT440-LARP7 antibody, we performed a peptide competition assay in which the pT440-LARP7 antibody (produced by Abclonal) was first blocked with mock peptide, pT440-LARP7 peptide, or non-pT440-LARP7 peptide and then used for WB detection.

### *In vitro* ubiquitination assay

*In vitro* ubiquitination assays were performed as previously described (Eakin et al., 2007). Briefly, purified His-LARP7 was mixed in an Eppendorf tube with 100 nM E1 (Boston Biochem), 400 ng of E2 (UbcH5c, Millipore, 23-035), 8 μM BRCA1/BARD1 RING domain, 2 μg of His-ubiquitin, 5 mM MgCl_2_, 0.5 mM DTT, 2 mM ATP, and 2 mM NaF in reaction buffer (50 mM Tris-Cl pH 7.5, 150 mM NaCl) and incubated in a 37°C water bath for 90 min. The reaction was stopped by boiling in 2x Laemmli loading buffer (63 mM Tris-HCl pH 6.8, 2% SDS, 20% glycerol and 0.01% bromophenol blue). WB analysis was used to detect the resulting ubiquitinated LARP7 with a LARP7-specific antibody.

### *In vivo* ubiquitination assay

Full-length LARP7, BRCA1 and BARD1 sequences together with His-ubiquitin were transfected into 293T cells for 48 hours, and 20 μM MG132 (Sigma-Aldrich, M7449) was added 8 hours before harvest. After saving aliquots of cells (≈10%) for quantification of protein expression, the remaining cells were lysed with buffer A (6 M guanidine HCl, 0.1 M Na_2_HPO_4_/NaH_2_PO_4_, and 5 mM imidazole) with brief sonication, and 10 mg of cell lysate was incubated with 50 µl of TALON Metal Affinity Resin and rotated at 4°C overnight. The beads were sequentially washed once with buffer A, twice with buffer B (1.5 M guanidine HCl, 25 mM Na_2_HPO_4_/NaH_2_PO_4_, 20 mM Tris-Cl pH 6.8, and 10 mM imidazole) and three times with buffer T1 (25 mM Tris-Cl pH 6.8 and 15 mM imidazole) and boiled with 100 µl of 2x Laemmli loading buffer containing 200 mM imidazole to release the ubiquitinated protein. The proteins were then detected with WB analysis.

### Bimolecular fluorescence complementation assay

BiFC was performed as previously described to detect LARP7-BARD1 interactions (Shyu et al., 2006). Briefly, 293T cells were seeded into 6-well plates for 24 hours after cotransfection with pBiFC-bFos-VC155 and pBiFC-bJun-VN173 or pBiFC-BARD1-VC155 and pBiFC-LARP7-VN173. The cells were then analyzed with a Nikon A1Si confocal microscope or with FACS for fluorescence detection.

### Coimmunoprecipitation

Co-IP experiments were performed as previously described (Chen et al., 2017). To prepare beads covalently conjugated with antibody, 10 μl of agarose protein G beads (GE Healthcare, 17-0618-01) were washed with PBS three times, incubated with 1 μg of antibody at 4°C for 6 hours and then crosslinked with 1 ml of PBS containing 25 mM dimethyl pimelimidate dihydrochloride (Thermo Fisher) for 1 hour at room temperature. The crosslinking reaction was terminated by washing the beads with 1 ml of 50 mM Tris-HCl (pH 7.4) for 5 min and washing them twice with PBS containing protease and phosphatase inhibitors.

The cell pellets were incubated with buffer A (10 mM HEPES pH 7.9, 10 mM KCl, 1.5 mM MgCl2, and 0.5 mM DTT) at 4°C for 10 min and centrifuged at 4000 rpm for 5 min. The precipitated nuclear pellets were lysed in high-salt buffer B for 10 min. The nuclear extracts were sonicated briefly and diluted with two volumes of dilution buffer (20 mM HEPES pH 7.9 and 0.2 mM EDTA plus protease and phosphatase inhibitors) before incubating with the corresponding antibody-conjugated beads. For endogenous immunoprecipitation, cell lysates (1-5 mg) were incubated at 4°C overnight with specific antibody-crosslinked protein G beads or normal rabbit isotype IgG-crosslinked beads as controls (Table S1). After rinsing with PBS three times, the pulldown proteins were eluted with 50 µl of elution buffer (50 mM glycine pH 2.5) for 5 min at room temperature, neutralized with 50 μl of 1 M Tris-Cl (pH 7.4) and then boiled with 20 μl of 6x Laemmli buffer.

To precipitate ectopically expressed proteins, pHAGE-FLAG-LARP7 was transfected into 293T cells together with pHAGE-HA-BRCA1, pHAGE-HA-BRCT (BRCA1), pHAGE-HA-BARD1 or pHAGE-HA-BRCT (BARD1). The cells were incubated with 20 μM MG132 for 8 hours and irradiated with various doses of IR. Cell lysate (1-5 mg) was immunoprecipitated with Flag antibody-conjugated M2 agarose beads (Sigma-Aldrich) at 4°C overnight, and after three washes with PBS, the precipitated protein was collected with 2x Laemmli loading buffer.

### Protein half-life assay

HeLa cells (1×10^6^) were treated with 25 μg/ml cycloheximide (CHX; M4879, Abmole) 15 min prior to IR or CDDP administration. The cells were lysed immediately with buffer B containing protease inhibitors at the indicated time points, and then the proteins were detected with WB analysis.

### Immunohistochemistry

Immunohistochemistry was performed on breast cancer tissue as previously described (Zhang et al., 2009). Paraffin-embedded breast cancer tissue arrays (BR804b and BR1321) purchased from Alenabio (US Biomax) were heated at 60°C for 30 min, deparaffinized in xylene and rehydrated in an ethanol gradient. Antigen retrieval was performed in citrate retrieval buffer (10 mM sodium citrate pH 6.0) at 95°C for 20 min. After washing with PBS, 3% H_2_O_2_ in methanol was applied to quench endogenous peroxidase. IHC was performed with an R.T.U. VECTASTAIN Universal ABC Kit (Vector Laboratories, PK-7200) following the manufacturer’s instructions. Briefly, the sections were color-developed with ImmPACT DAB Substrate (Vector Laboratories, SK-4105) and counterstained with hematoxylin after sequential incubation with primary antibodies (1:100 to 1:200 dilutions, Table S1), biotinylated secondary antibody and R.T.U. VECTASTAIN ABC Reagent. A Nikon DS-Ri2 microscope was used to image the stained sections. LARP7 and BRCA1 signals were scored as negative (0), weak (1), intermediate (2) or strong (3) for Wilcoxon rank-sum tests or were classified into low (negative and weak) or high (intermediate and strong) groups for Fisher’s exact tests.

### Immunofluorescence

Cells were seeded on glass coverslips for 24 hours, collected at the times indicated after X-ray IR (10 Gy) and fixed with 4% paraformaldehyde for 10 min. After blocking with buffer (5% normal donkey serum, 1% BSA and 0.3% Triton X-100 in PBS) for 30 min, the coverslips were incubated sequentially with primary antibodies (Table S1) at 1:100 to 1:200 dilutions overnight at 4°C, Alexa Fluor 488- and 555-conjugated secondary antibodies (1:200, Thermo Fisher) for 1 hour at room temperature and 4,6-diamidino-2-phenylindole (DAPI) or Hoechst 33342 (Invitrogen) nuclear dye for 10 min at room temperature. The coverslips were mounted on microscope slides using ProLong Gold Antifade Reagent (Invitrogen) and imaged under a Nikon A1Si confocal microscope.

### RNA-seq

RNA-seq was performed as previously described (Chen et al., 2017). Total RNA was purified by using an RNeasy Miniprep Kit (Qiagen) with an on-column DNase I digestion protocol to eliminate residual DNA. RNA purity was monitored with a NanoPhotometer® spectrophotometer (Implen). mRNA was purified from 1.5 µg of total RNA using poly-T oligo magnetic beads, and libraries were constructed with an NEBNext® Ultra^TM^ RNA Library Prep Kit for Illumina (NEB). An AMPure XP system was used for DNA purification, and library quality was assessed with an Agilent Bioanalyzer 2100 system. Next-generation sequencing was performed on an Illumina HiSeq 4000 platform by Novogene, and 150 bp paired-end reads were generated.

### Cell cycle analysis

Cells were collected and fixed in ice-cold 70% ethanol overnight at 4°C, permeabilized with 0.25% Triton X-100 in PBS, incubated with an anti-pH3 antibody (Table S1) for 2 hours at room temperature, and incubated with an Alexa Fluor 488-conjugated secondary antibody for 1 hour at room temperature in the dark. DNA was stained with 20 μg/ml propidium iodide (PI) in the presence of 100 μg/ml RNase A. The stained cells were detected using an LSRFortessa flow cytometer (BD Biosciences), and the data were analyzed using FlowJo software.

### Chromosomal abnormalities

LARP7^+/+^ MEFs, LARP7^-/-^ MEFs or LARP7^-/-^ MEFs with recovered LARP7 were irradiated with X-rays at dose of 2 Gy. Twenty-four hours later, the MEFs were exposed to colcemid (0.1 μg/ml, Roche) for 6 hours, further subjected to hypotonic treatment (75 mM KCl), and fixed in a methanol and acetic acid mixture (3:1). Chromatin was stained in Giemsa solution (Solarbio) and visualized for structural abnormalities under a ZEISS microscope (MetaSystems). At least 50 metaphase spreads were analyzed in each sample.

### Homologous repair assay

A homologous repair assay was performed as previously described (Peng et al., 2009). Briefly, stable DR-GFP cell lines (U2OS DR-GFP was a gift from Dr. Shiaw-Yih Lin and ESC DR-GFP was a gift from Dr. Anyong Xie) were transfected with control or LARP7 siRNAs or overexpression vectors. After 24 hours, the cells were further transfected with the I-SceI expression vector pCBASce. GFP-positive cells were detected using an LSRFortessa flow cytometer (BD Biosciences) two days after transfection.

### Computational analysis

#### RNA-seq analysis

Raw reads were filtered using Cutadapt (https://cutadapt.readthedocs.io/en/stable/) and checked with FastQC. Filtered reads were mapped to GRCh37 using STAR with GENCODE19 annotations. The mapped reads for each gene were counted using GFOLD (https://bitbucket.org/feeldead/gfold) with the default settings, and the expression changes were quantified using GFOLD diff, the calculations of which are based on the posterior distribution of the log fold changes. A threshold of |GFOLD|≥0.585 was set to select differentially expressed genes. Differentially expressed genes in MEFs, HCC1937 cells and 231 cells were imported into Ingenuity Pathway Analysis (IPA, Qiagen), and a core analysis was used to reveal dysregulated pathways in LARP7-knockout or LARP7-knockdown cell lines. The expression of select genes related to the cell cycle were normalized, grouped with hierarchical clustering and plotted in R with the pheatmap package.

#### Cancer survival analysis

KMplot (http://kmplot.com/analysis/) was used to analyze the prognostic value of LARP7, BRCA1 and BRCA2. Patients were divided into two groups with high (top 25%) and low (bottom 25%) expression. Kaplan-Meier survival plots were used to evaluate patient RFS and to calculate the hazard ratios and average survival in months. Significant differences between two groups were determined with logrank tests.

---

**Figure S1.**
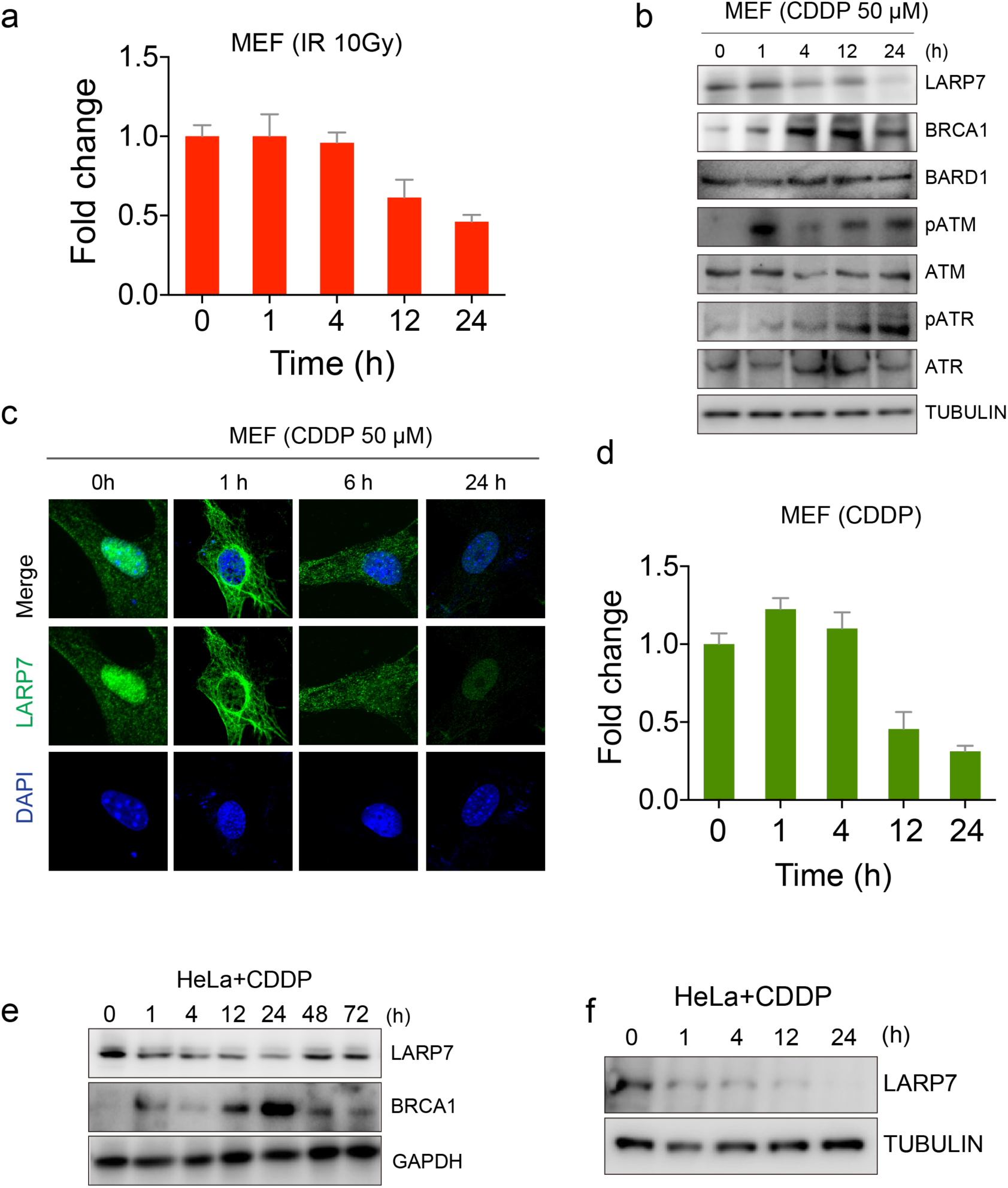
LARP7 was attenuated and extranuclearly shuttled during the DDR. a, IR treatment lead to the decrease of 7SK RNA at 12 and 24 hours in MEF cells. b, The chemotherapeutic drug CDDP suppressed LARP7 expression in MEFs. c, Immunofluorescence staining illustrating the downregulation and extranuclear movement of LARP7 after CDDP treatment. d, CDDP resulted in a decreasing of 7SK RNA in MEF cells. e, CDDP repressed LARP7 expression for up to 24 hours in HeLa cells, and this effect was coordinated with BRCA1 upregulation. f, LARP7 downregulation upon CDDP treatment was validated by WB analysis with a LARP7 antibody from a different manufacturer.

**Figure S2.**
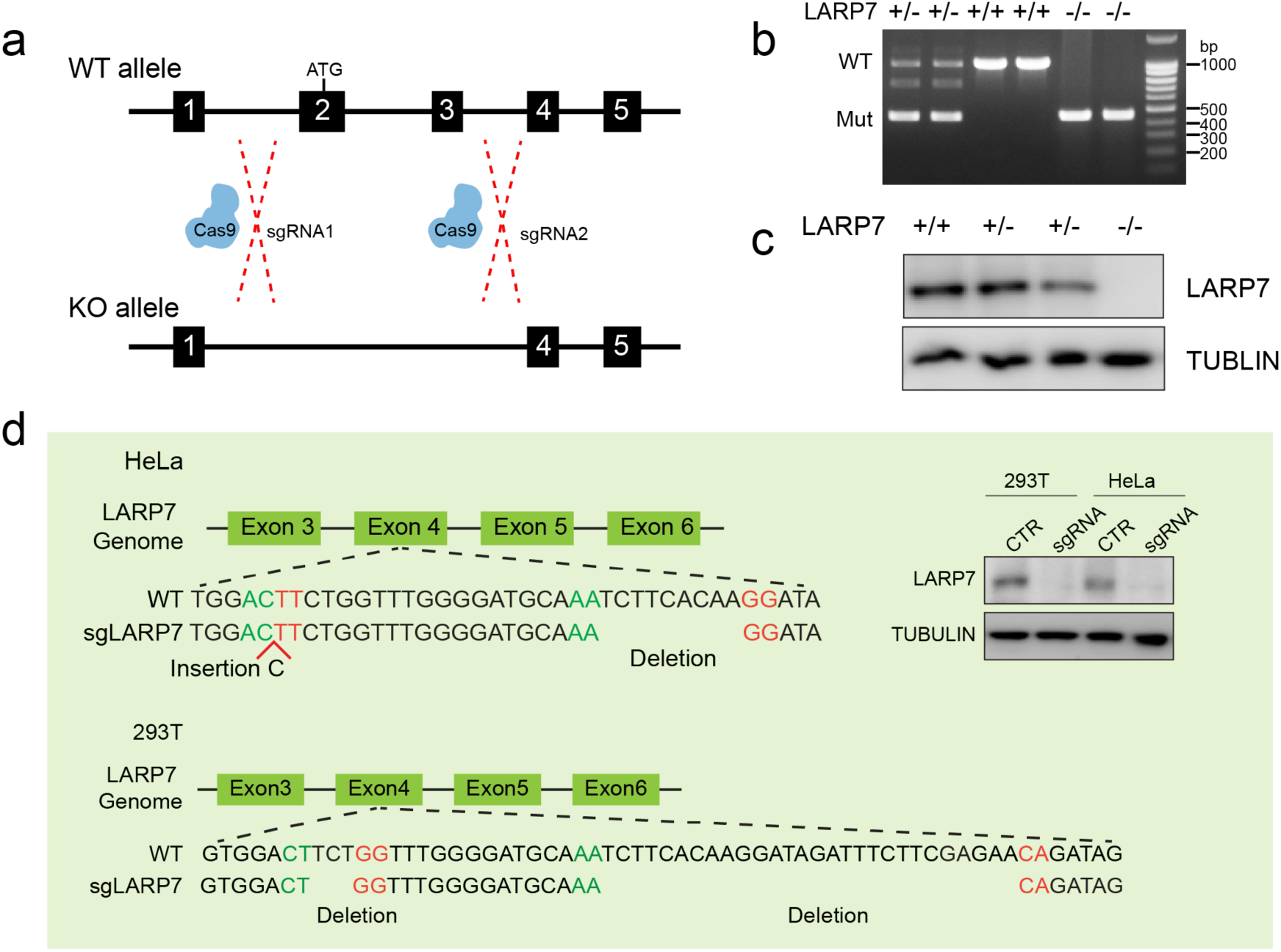
Generation of LARP7 knockout MEF and HeLa cells. a, Targeted strategy for development of LARP7 conventional knockout (KO) mice. b, Genotyping gel for LARP7^+/+^, LARP7^+/-^ and LARP7^-/-^ mice. c, WB validating that there was no LARP7 protein expression in LARP7^-/-^ MEFs. d, Validation of the LARP7^-/-^ HeLa and 293T cell lines generated by CRISPR. The left side of the panel shows the genotype of the LARP7^-/-^ HeLa and 293T cell lines. The right side of the panel shows a WB verifying LARP7 protein depletion.

**Figure S3.**
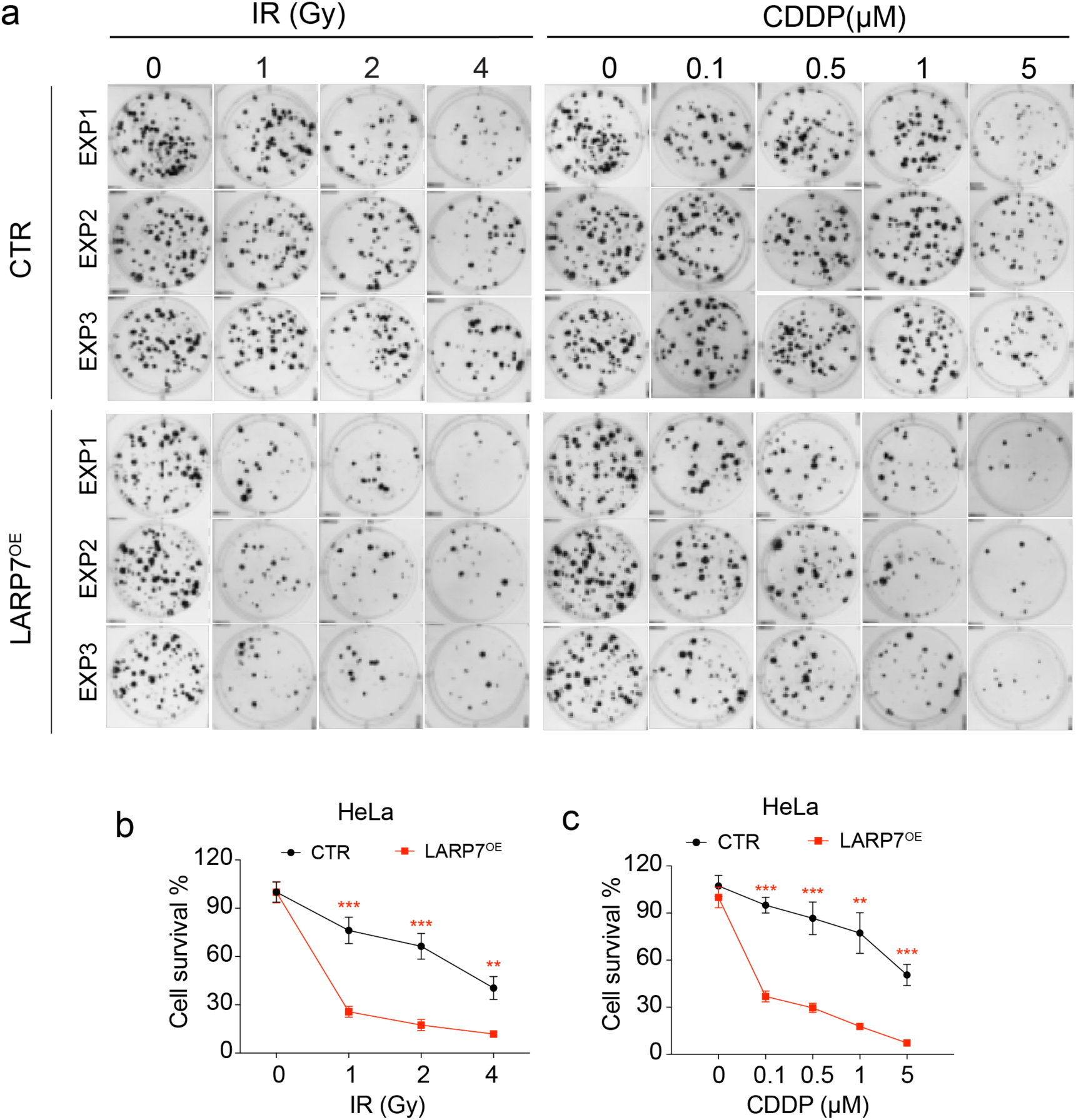
LARP7 decreased HeLa cell survival upon DNA damage. a-c, Clone formation assay of wild-type and LARP7-overexpressing HeLa cells treated with IR (a, b) and CDDP (a, c). Mean±SD, Student’s t test, n=3, *: p<0.05, **: p<0.01.

**Figure S4.**
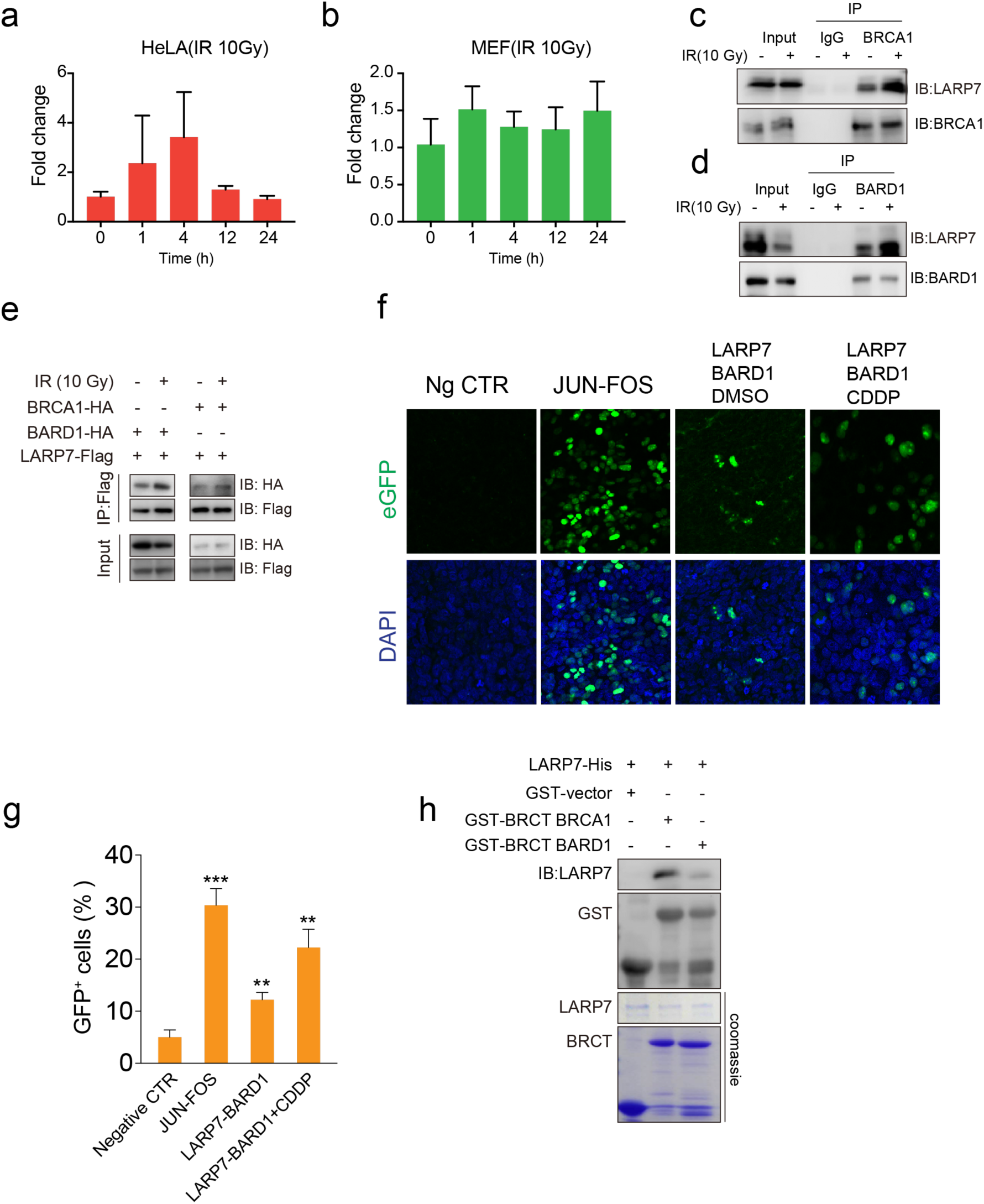
The BRCA1/BARD1 complex ubiquitinated LARP7. a-b, IR did not suppress LARP7 mRNA transcription in HeLa cells (a) and MEFs (b), as detected by RT-qPCR. Data: mean±SD, Student’s t test, n=3. c-d, Reciprocal IP demonstrated that IR increased the interaction between LARP7 and wild-type BRCA1/BARD1 in 293T cells. e, A Co-IP experiment indicated that IR increased the interaction between LARP7 and BRCA1/BARD1. f-g, A BiFC assay showed that CDDP enhanced the interaction between LARP7 and BARD1. Green cells indicate cells with positive interactions. The JUN-FOS interaction was used as a positive control. Student’s t test, n=3, *: p<0.05, **: p<0.01. h, A GST pulldown assay showed that the BRCT domain of BARD1 had much lower affinity for LARP7 than the BRCT domain of BRCA1.

**Figure S5.**
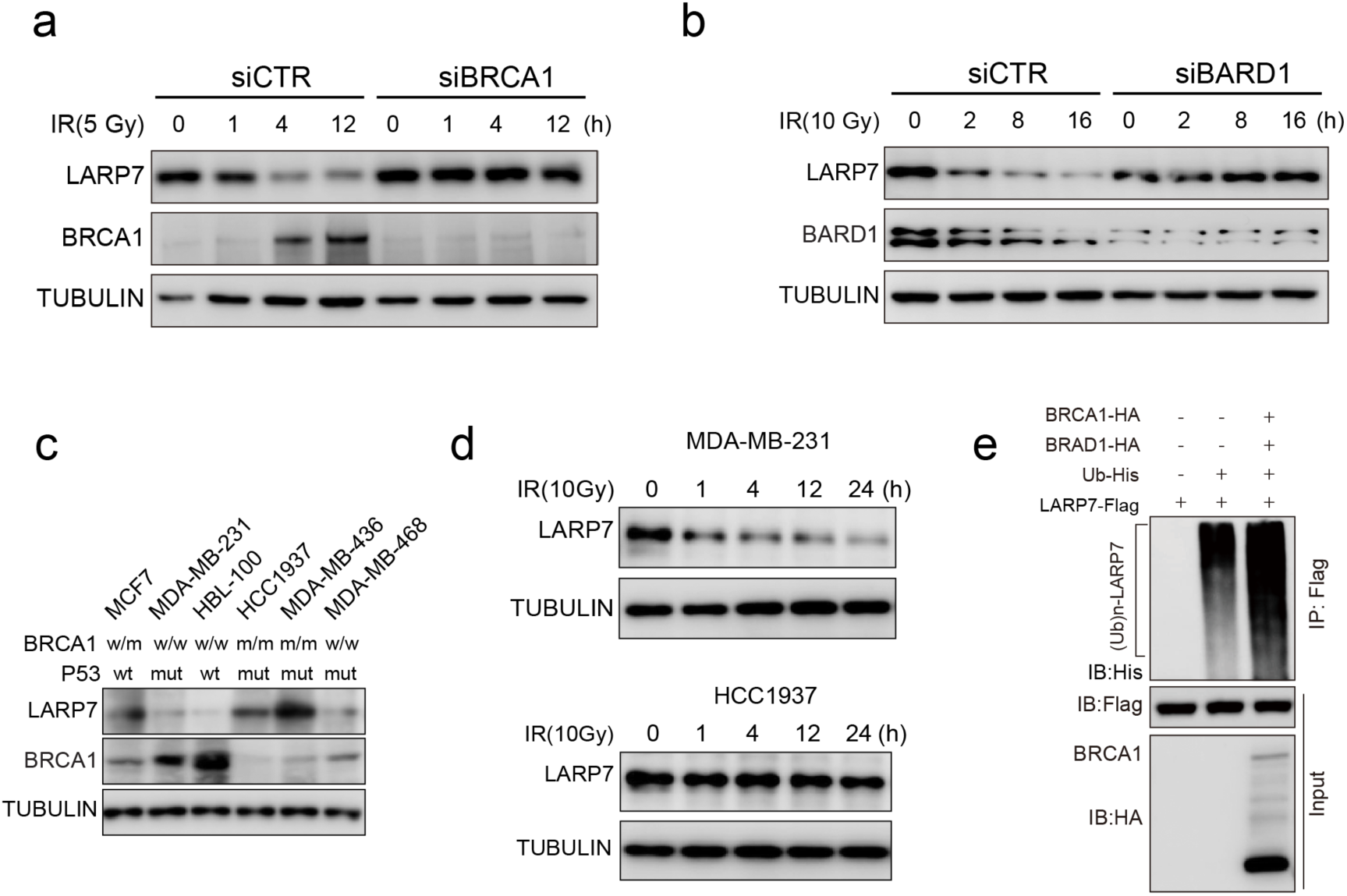
BRCA1/BARD1 ubiquitinated LARP7. a, BRCA1 knockdown attenuated IR-induced LARP7 degradation. b, BARD1 knockdown attenuated IR-induced LARP7 degradation. c, LARP7 expression in variable breast cancer cell lines. w: wild-type, m: BRCA1 mutant, mut: P53 mutant. MDA-MB-436 and HCC1937 cells have a homozygous BRCA1 mutation; MCF7 cells have a heterozygous mutation. d, IR induced LARP7 downregulation in MDA-MB-231 cells but not in HCC1937 cells. e, Ectopic expression of BRCA1 and BARD1 augmented LARP7 ubiquitination in 293T cells.

**Figure S6.**
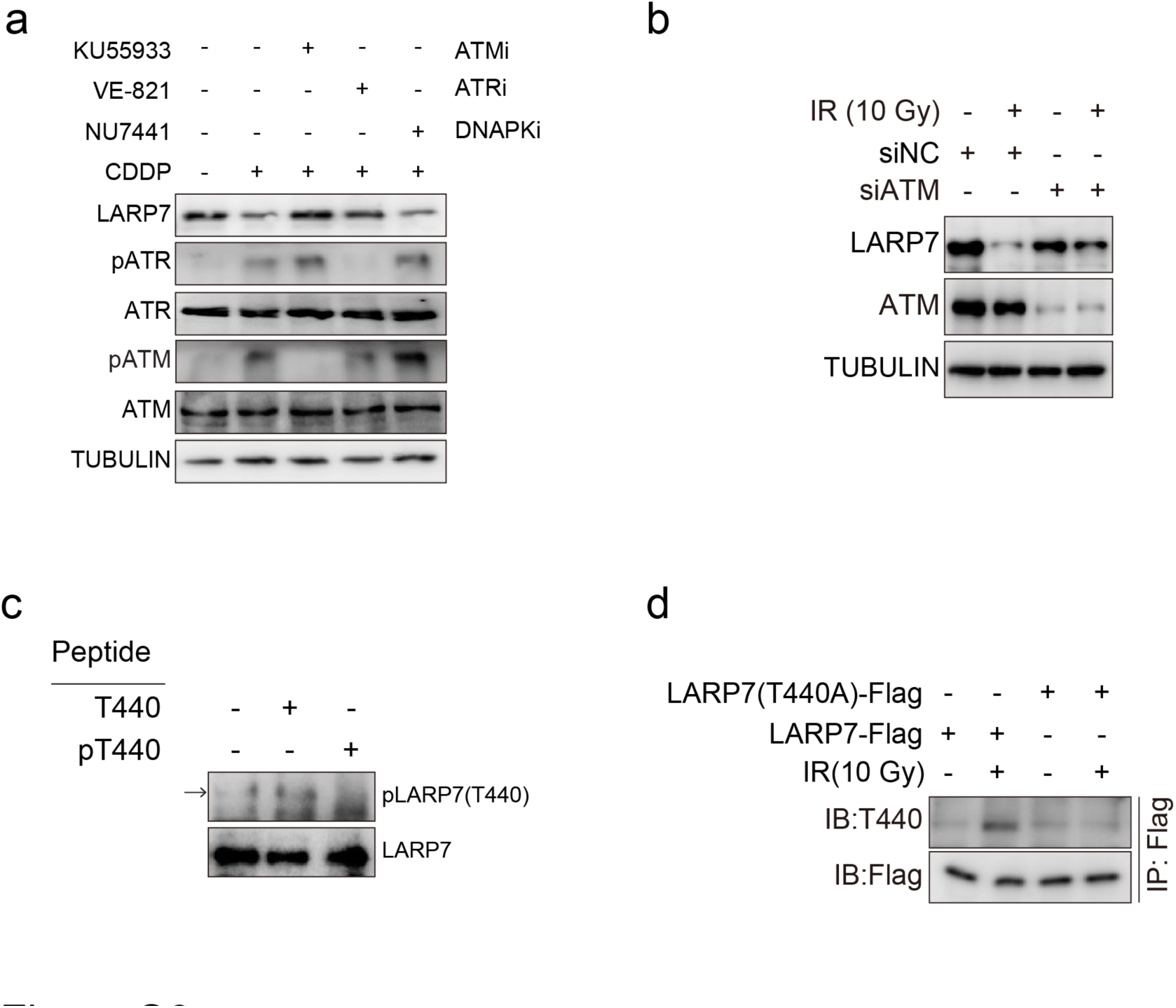
ATM-mediated T440 phosphorylation facilitated LARP7 interactions with BRCA1. a, ATM and ATR inhibitors (ATMI and ATRI, respectively; 10 μM), but not a DNAPK inhibitor, attenuated CDDP-induced LARP7 degradation. b, ATM siRNA attenuated IR-induced LARP7 downregulation. c, Preincubation with a T440 phosphorylation peptide abolished the recognition of LARP7 by the phosphorylation antibody, indicating its specificity. d, The T440A mutation abolished IR-induced T440 phosphorylation.

**Figure S7.**
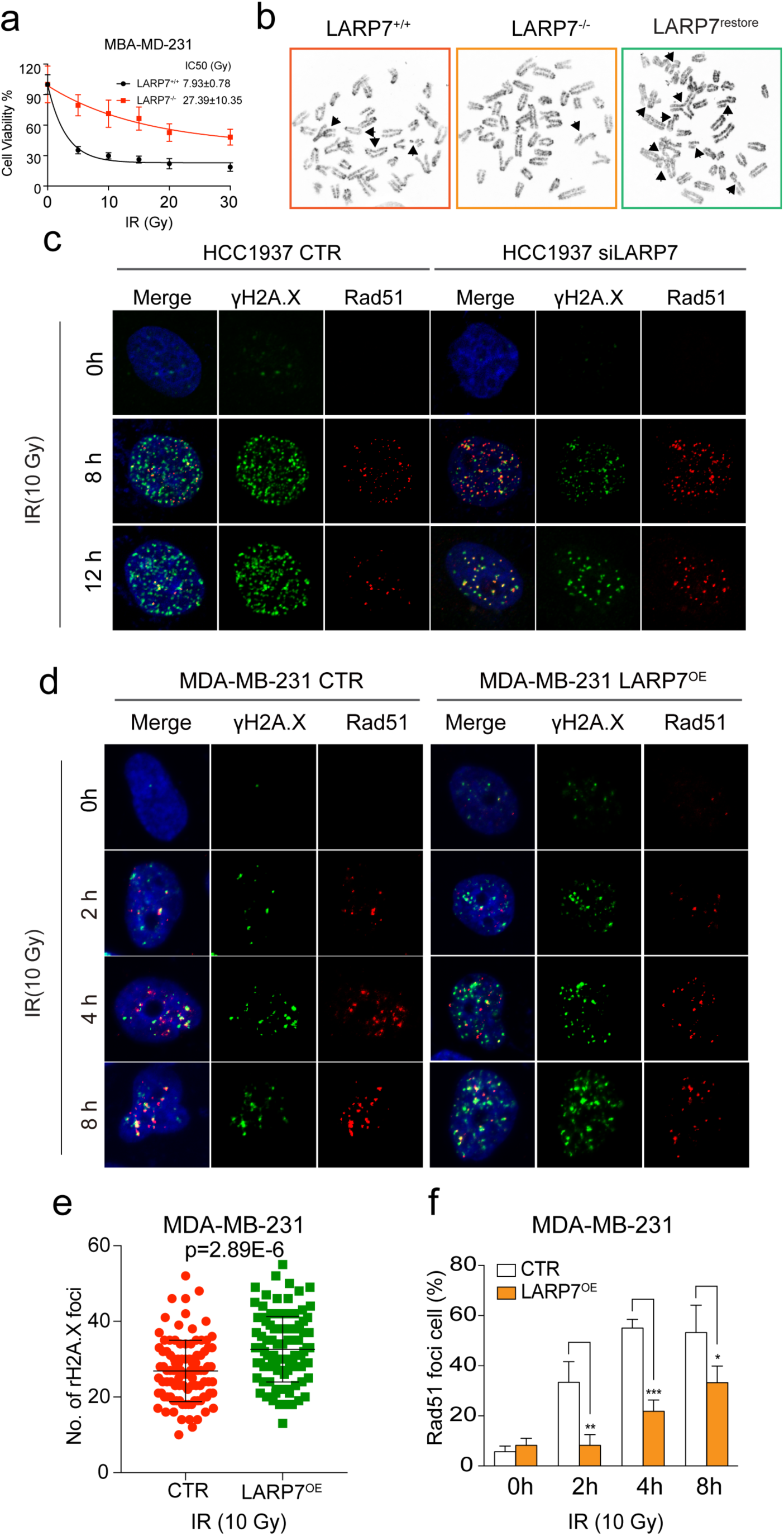
LARP7 attenuated RAD51 recruitment upon DNA damage. a, Cell survival curve of MDA-MB-231 cells. LARP7 depletion increased cell survival of IR-treated MDA-MB-231 cells. Data: mean±SD, Regression: one-phase decay model. b. Representative images of chromatin abnormalities. c, Images of Rad51 and γH2A.X foci in HCC1937 cells corresponding to Fig. 4g-h. d, Images of Rad51 and γH2A.X foci in MDA-MB-231 cells corresponding to sFig. 6e-f. e, Statistical summary of the number of γH2A.X foci per cell at 4 hours (MDA-MB-231 cells) after IR. Data: mean±SD, Student’s t test, n=100 cells. f, Percentage of RAD51 focus-positive MDA-MB-231 cells. LARP7 overexpression decreased the percentages of RAD51 foci-positive cells at the three tested times. Student’s t test, n=300-400, *: p<0.05, **: p<0.01, ***: p<0.001.

**Figure S8.**
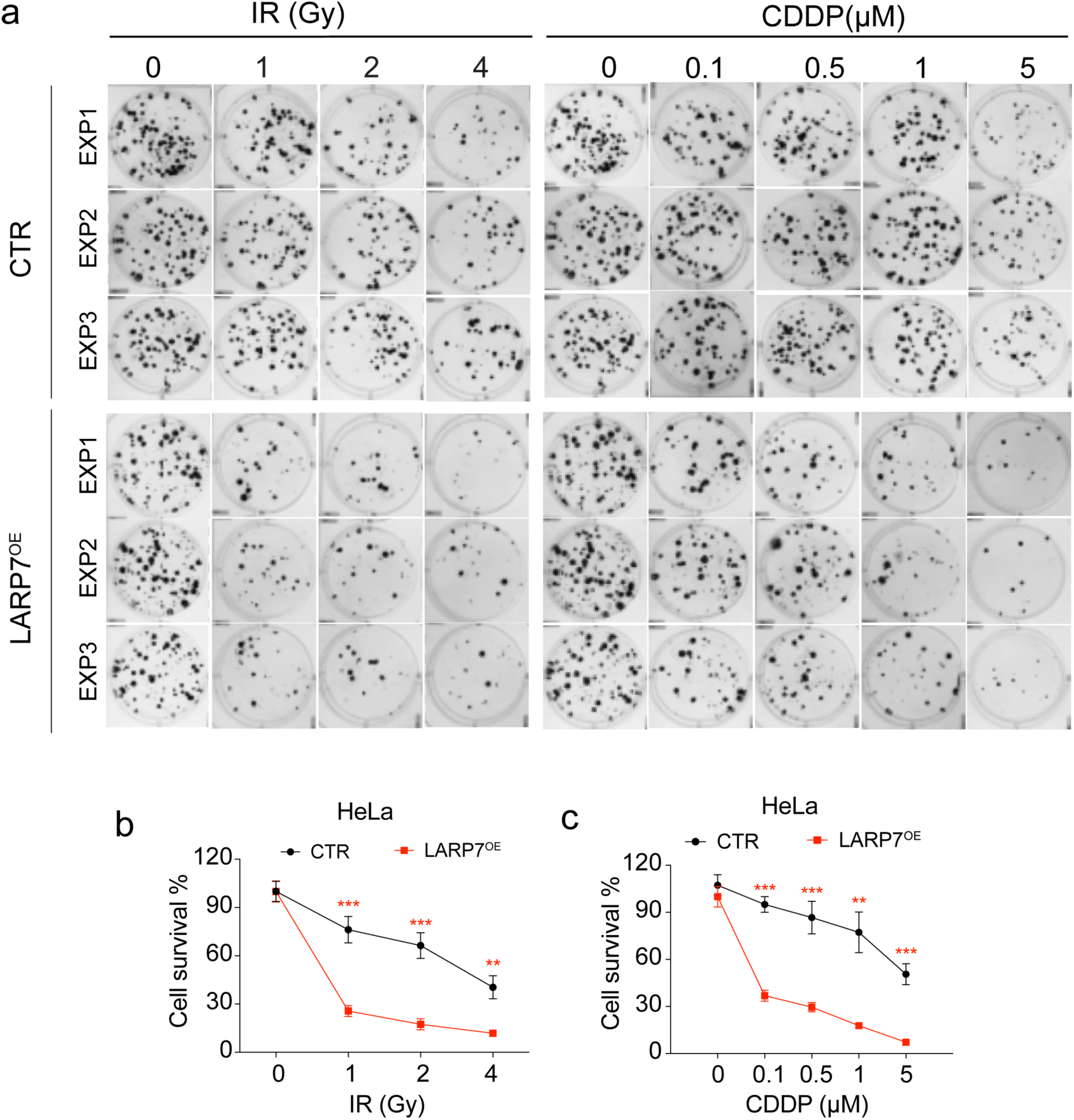
LARP7 increased the susceptibility of breast cancer’s to chemo- and radiotherapy. a LARP7 overexpression increased the sensitivity of MDA-MB-231 cells to IR. Data: mean±SD, Regression: one-phase decay model. b-c, Clonogenic assay showed that LARP7 increased the susceptibility of MDA-MB-231 cells to IR LARP7. b, Quantification of survival clones. Mean±SD, Student’s t test, n=3, *: p<0.05, **: p<0.01. c, Images of survival MDA-MB-231 clones after IR.

**Figure S9.**
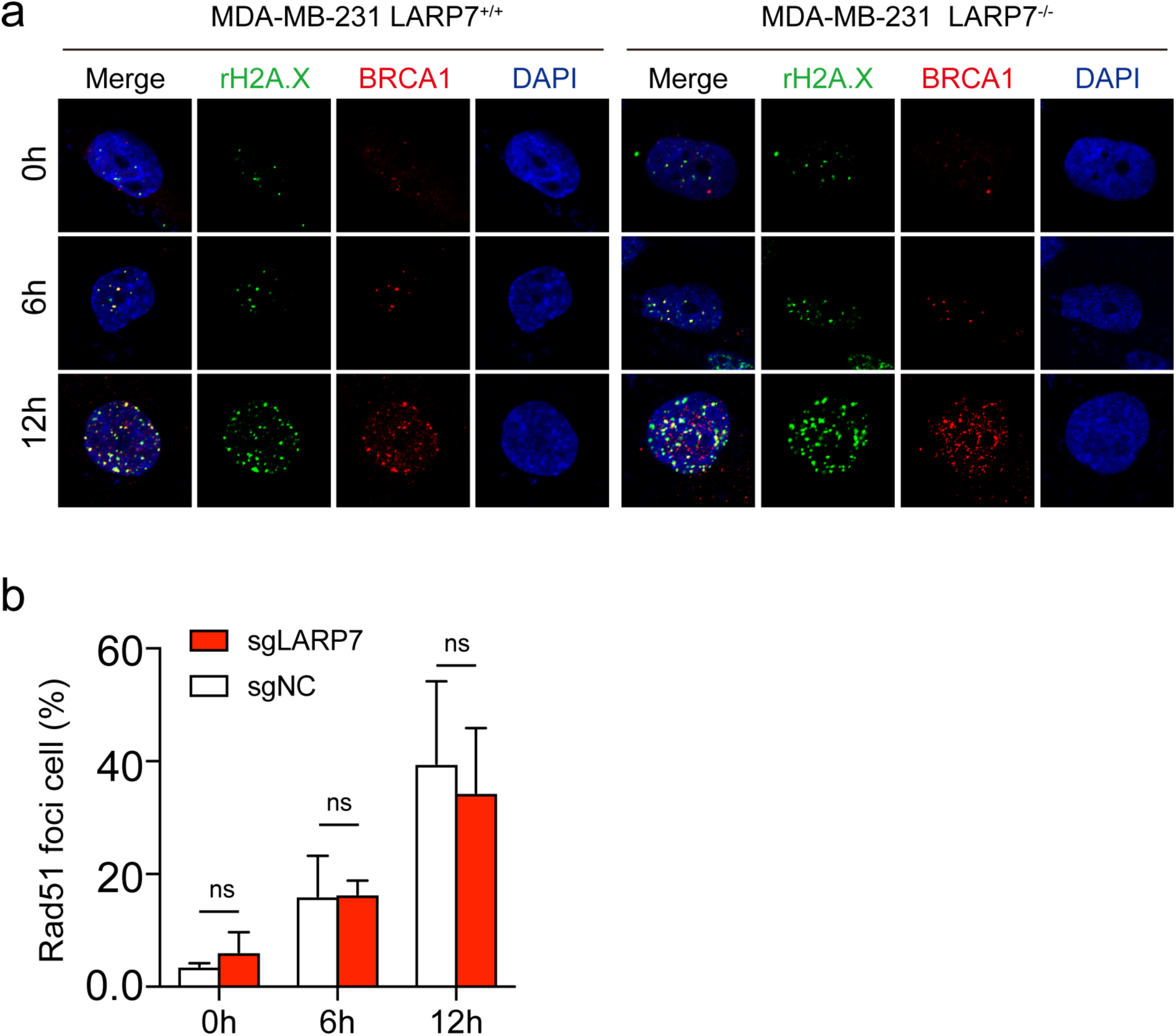
LARP7 depletion had no effect on BRCA1 foci formation. a, Cell survival curve of MDA-MB-231 cells. LARP7 depletion increased cell survival of IR-treated MDA-MB-231 cells. Data: mean±SD, Regression: one-phase decay model. b, Representative images of chromatin abnormalities corresponding to Figure 5a. c, Representative images of BRCA1 foci upon IR (10 Gy). d, Statistical summary of BRCA1 foci. Data: mean±SD, student’s t test, ns: no significance, n=150 cells.

**Figure S10.**
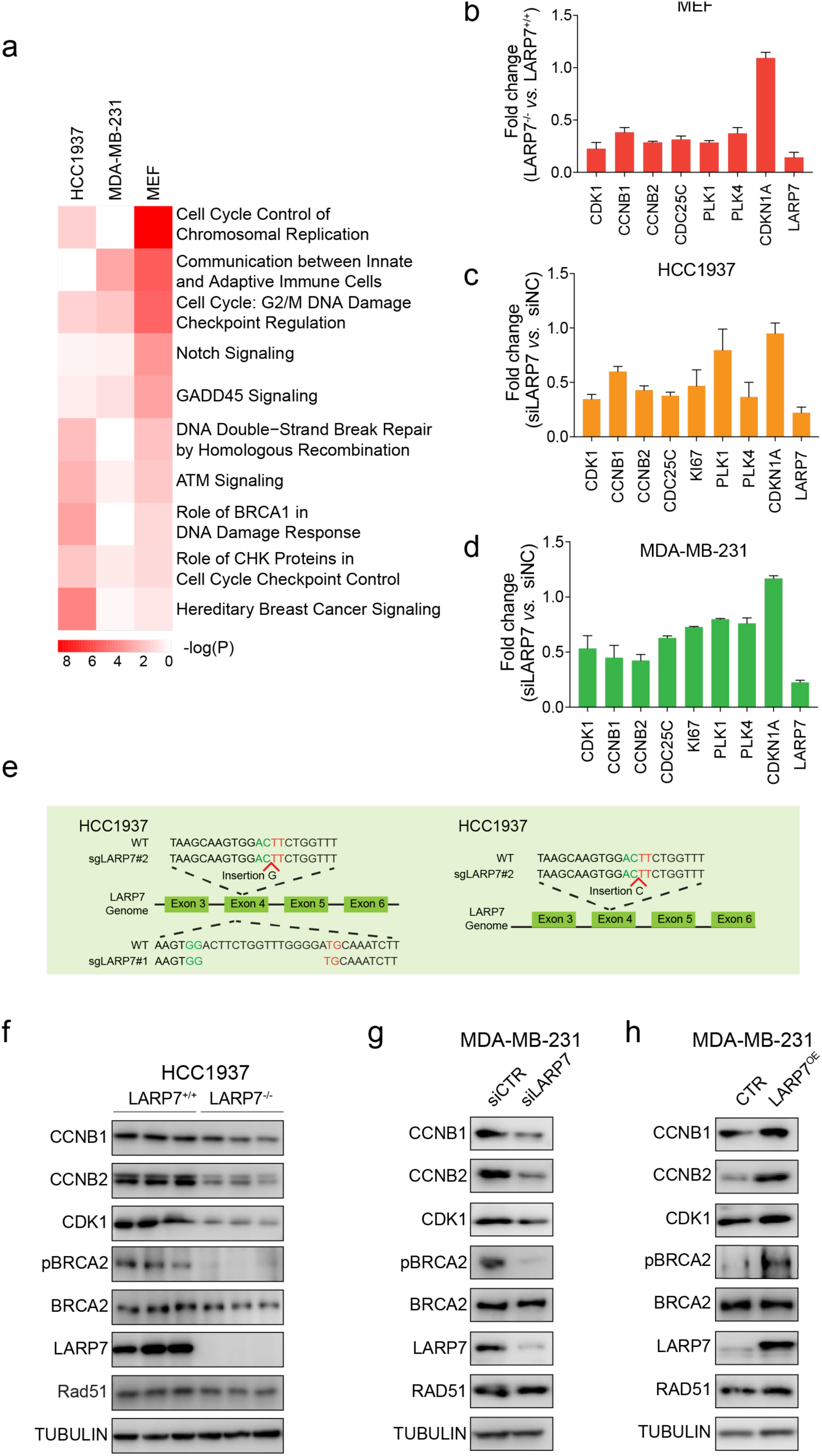
LARP7 enhanced the expression of CCNB1, CCNB2 and CDK1. a, Enriched biological terms for the differentially expressed genes in LARP7-depleted cell lines as determined by IPA analysis. b-d, LARP7 depletion attenuated cell cycle gene expression in MEFs (b), HCC1937 cells (c) and MDA-MB-231 cells (d), as revealed by RT-qPCR. Data: mean±SD, n=3-4. e, Validation of the three LARP7^-/-^ HCC1937 cell lines created with two independent sgRNAs. Sanger sequencing was used to genotype the HCC1937 cell lines. f, LARP7-knockout HCC1937 cells had lower protein levels of CCNB1, CCNB2, and CDK1 and lower phosphorylation of BRCA2 than wild-type cells. g-h, In MDA-MB-231 cells, knockdown of LARP7 decreased CCNB1, CCNB2, and CDK1 expression and BRCA2 phosphorylation (g), while overexpression increased these parameters (h).

**Figure S11.**
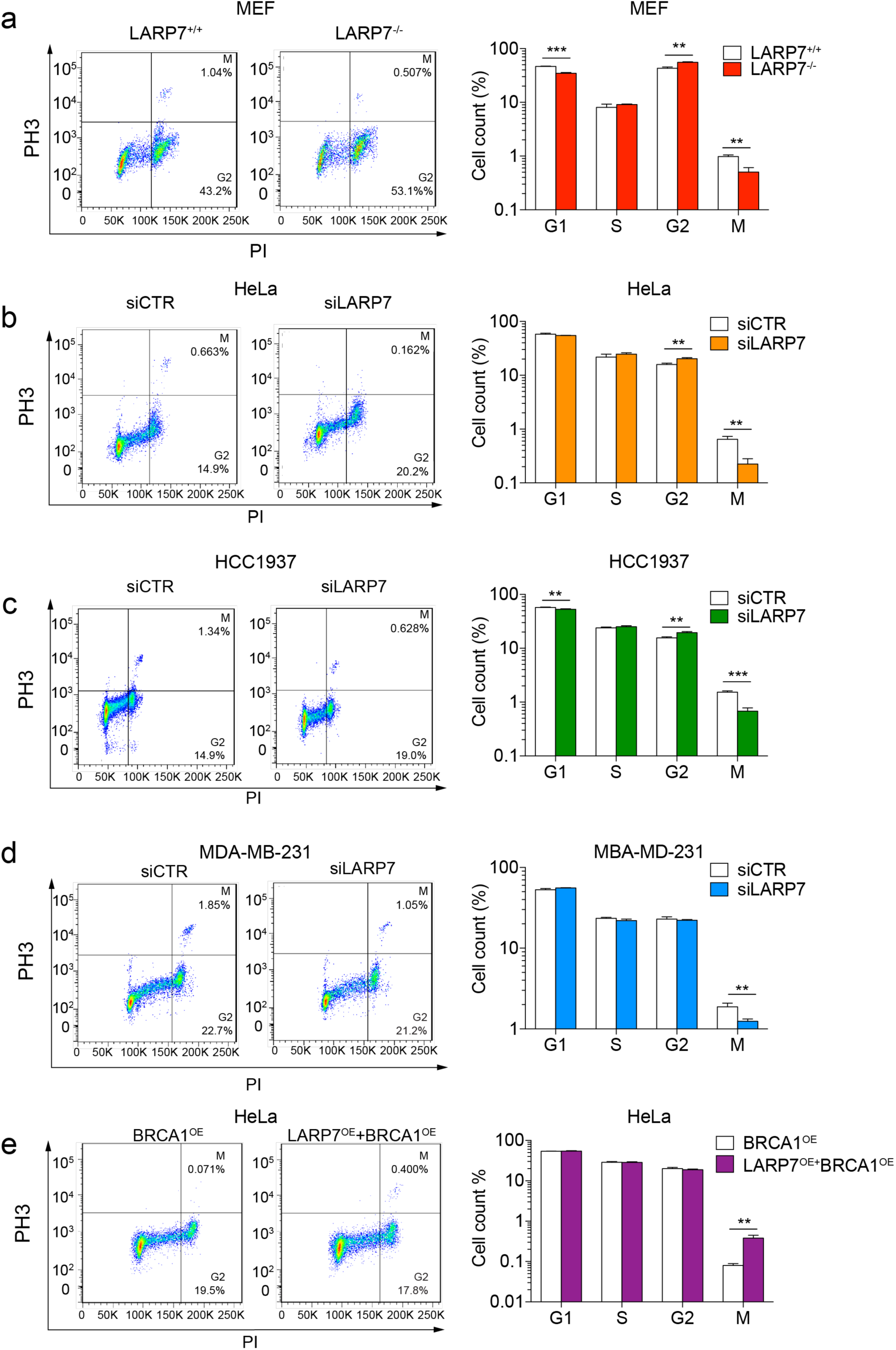
LARP7 promoted the G2/M cell cycle transition. a-c, LARP7 depletion induced G2/M cell cycle arrest in MEFs (a), HeLa cells (b), HCC1937 cells (c) and MDA-MB-231 cells (d). e, LARP7 overexpression enabled HeLa cells to overcome BRCA1-induced G2/M arrest. Left panels: FACS images of pH3 and PI double staining. Right plots: summaries of cell cycle distribution. Mean±SD, Student’s t test, n=3, *: p<0.05, **: p<0.01, ***: p<0.001.

**Figure S12.**
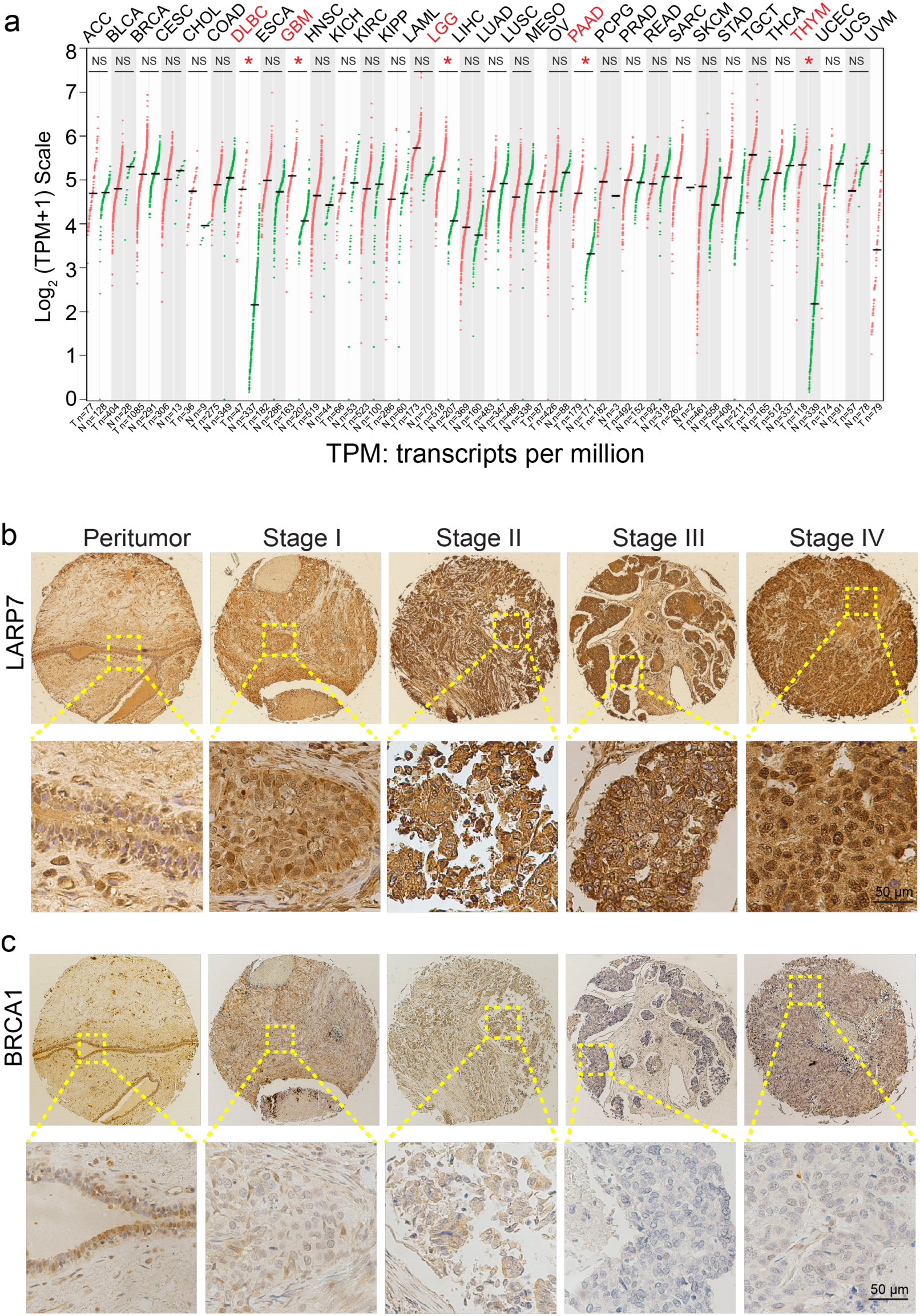
LARP7 was elevated in breast cancer tissues. a, LARP7 mRNA expression data for 33 types of human tumors obtained from the GEPIA database. Tumor samples of cancers highlighted with red have significantly different LARP7 mRNA expression than normal samples. b-c, Immunohistochemical staining showing LARP7 (b) and BRCA1(c) expression in breast cancers of various TNM stages corresponding to Fig. 7b.

**Figure S13.**
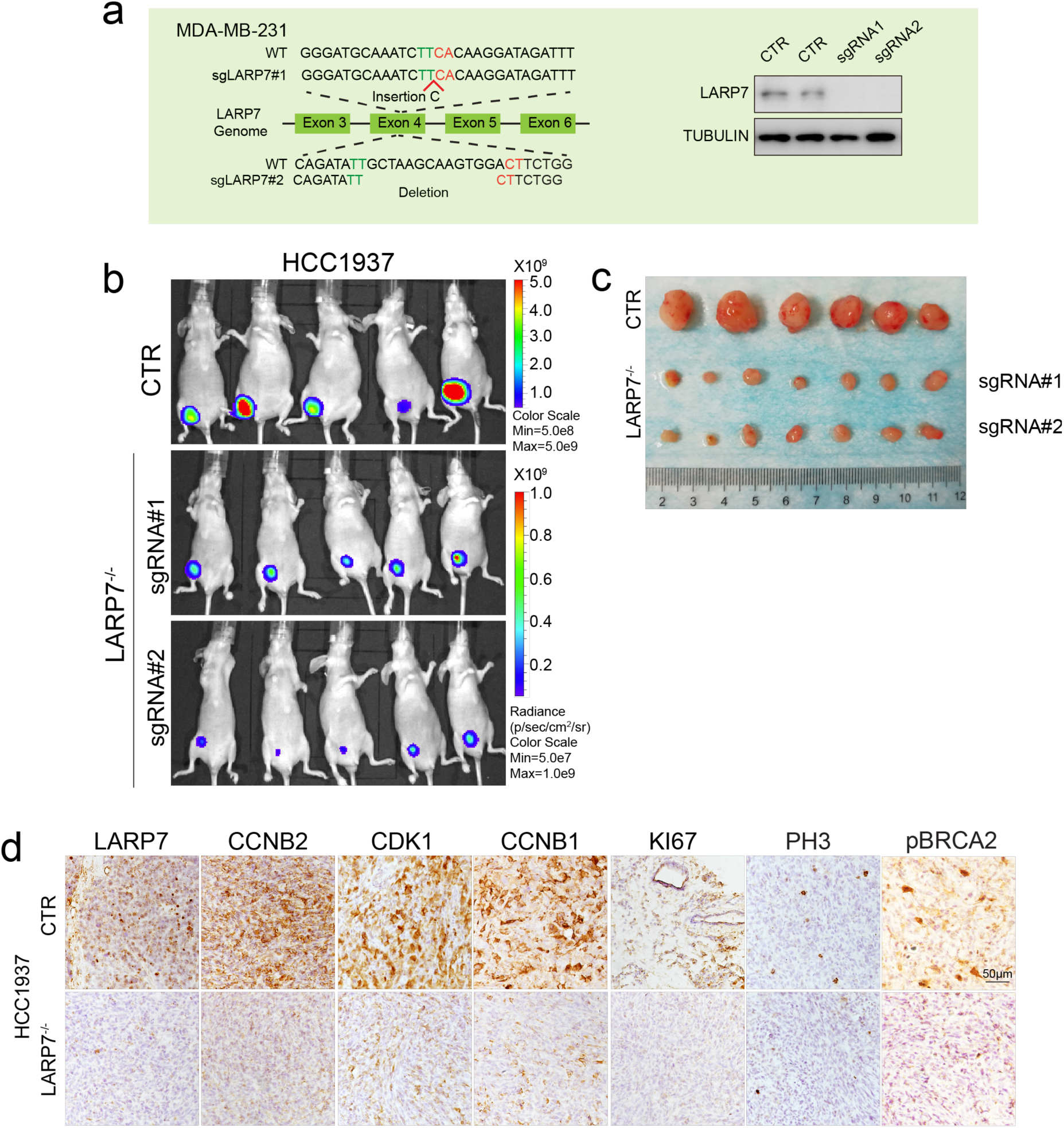
LARP7 promoted tumorigenesis. a, Validation of LARP7-knockout MDA-MB-231 cells. Genomic deletion generated with CRISPR was validated with Sanger sequencing (left panel), and protein depletion was validated with WB analysis (right panel). b-c. LARP7 depletion suppressed HCC1937 tumor growth. b, Luminescence images of HCC1937 xenograft tumors. c, Harvested tumor images corresponding to b. d, IHC images of wild-type and LARP7^-/-^ HCC1937 xenografts.

**Figure S14.**
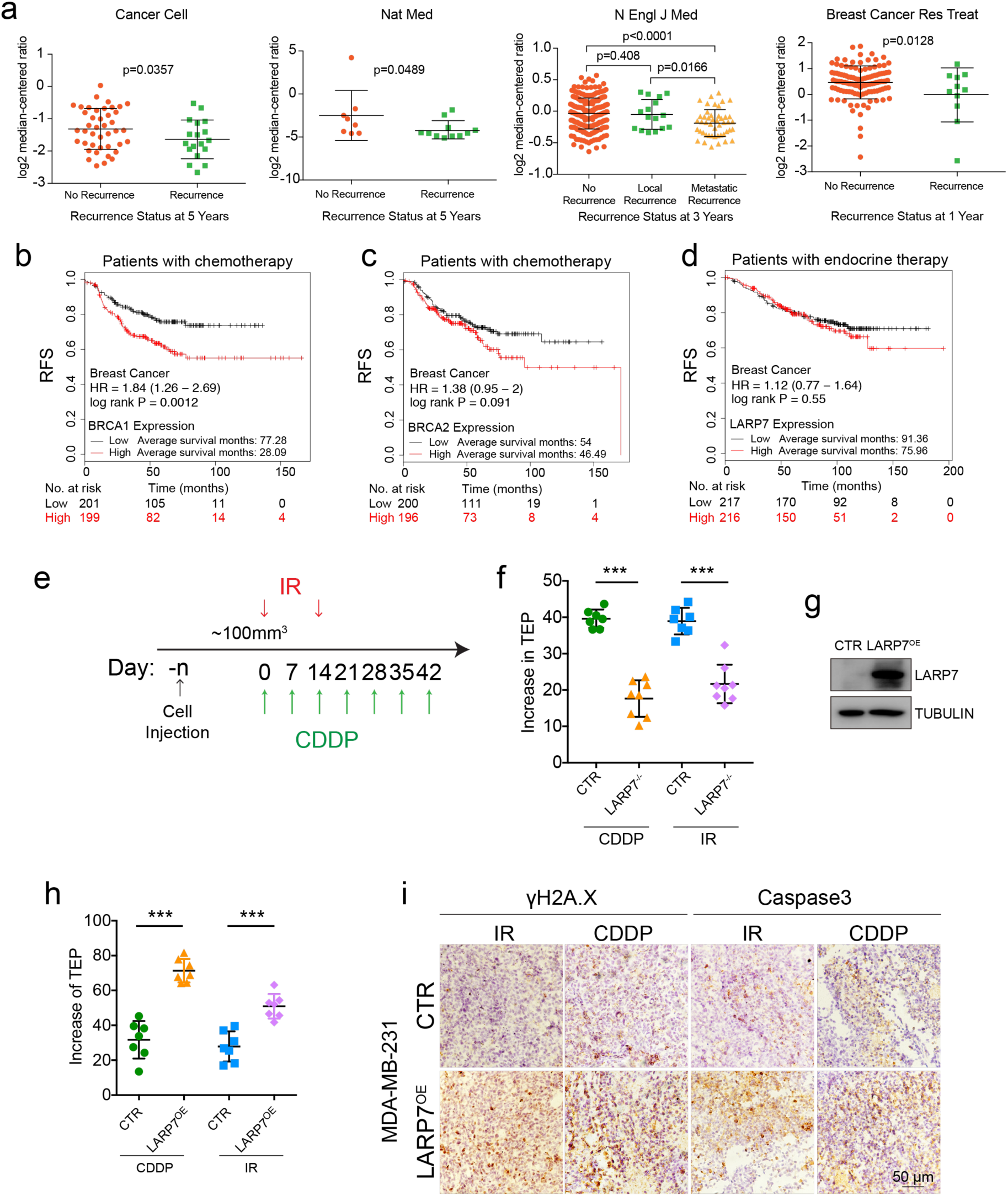
LARP7 increases RFS in breast cancer patients. a, LARP7 is downregulated in recurrent breast cancers. Student’s t test; the numbers of patients are indicated by the colored dots. The data were extracted from the Oncomine database. b-c, Patients with low levels of BRCA1 (b) and BRCA2 (c) have longer RFS. d, Patients with high LARP7 do not have better RFS after endocrine therapy. Logrank test; the numbers of patients that died at different times are labeled at the bottom of the plots. p<0.05 indicated significance. e, Schematic diagram of the chemo- and radiotherapy assay. e. Schematic diagram of the chemo- and radiotherapy assay. f, Increased TEP in wild-type and LARP7^-/-^ HCC1937 tumors upon treatment with CDDP or IR corresponding to Fig. 7c. The increases in TEP were smaller for LARP7^-/-^ tumors than for wild-type tumors. Data: mean±SD, n=7 mice for wild-type HCC1937 tumors, n=8 mice for LARP7^-/-^ HCC1937 tumors, unpaired Student’s t test. ***: p<0.001. g, WB validating LARP7 overexpression in injected LARP7^OE^ MDA-MB-231 cells. h, Increased TEP in wild-type and LARP7^OE^ MDA-MB-231 tumors upon treatment with CDDP or IR corresponding to Fig. 7d. The increases in TEP were greater in LARP7^OE^ tumor mice than in wild-type tumor mice. Data: mean±SD, n=7, unpaired Student’s t test. ***: p<0.001. i, IHC showing stronger γH2A.X and caspase-3 signals in IR- or CDDP-treated LARP7^OE^ tumors than in treated wild-type tumors.

